# Do You Want to Build a Genome? Benchmarking Hybrid Bacterial Genome Assembly Methods

**DOI:** 10.1101/2021.11.07.467652

**Authors:** Georgia L Breckell, Olin K Silander

## Abstract

Long read sequencing technologies now allow routine highly contiguous assembly of bacterial genomes. However, because of the lower accuracy of some long read data, it is often combined with short read data (e.g. Illumina), to improve assembly quality. There are a number of methods available for producing such hybrid assemblies. Here we use Illumina and Oxford Nanopore (ONT) data from 49 natural isolates of *Escherichia coli* to characterise differences in assembly accuracy for five assembly methods (Canu, Unicycler, Raven, Flye, and Redbean). We evaluate assembly accuracy using five metrics designed to measure structural accuracy and sequence accuracy (indel and substitution frequency). We assess structural accuracy by quantifying (1) the contiguity of chromosomes and plasmids; (2) the fraction of concordantly mapped Illumina reads withheld from the assembly; and (3) whether rRNA operons are correctly oriented. We assess indel and substitution frequency by quantifying (1) the fraction of open reading frames that appear truncated and (2) the number of variants that are called using Illumina reads only. Applying these assembly metrics to a large number of *E. coli* strains, we find that different assembly methods offer different advantages. In particular, we find that Unicycler assemblies have the highest sequence accuracy in non-repetitive regions, while Flye and Raven tend to be the most structurally accurate. In addition, we find that there are unidentified strain-specific characteristics that affect ONT consensus accuracy, despite individual reads having similar levels of accuracy. The differences in consensus accuracy of the ONT reads can preclude accurate assembly regardless of assembly method. These results provide quantitative insight into the best approaches for hybrid assembly of bacterial genomes and the expected levels of structural and sequence accuracy. They also show that there are intrinsic idiosyncratic strain-level differences that inhibit accurate long read bacterial genome assembly. However, we also show it is possible to diagnose problematic assemblies, even in the absence of ground truth, by comparing long-read first and short-read first assemblies.

**Author Notes:** All supporting data, code and protocols have been provided within the article or through supplementary data files. The supporting code is available from the GitHub repository https://github.com/GeorgiaBreckell/assembly_pipeline. nine supplementary figures and three supplementary tables are available with the online version of this article.

**Data summary:** Sequence data and genome assemblies for the natural isolates are available at https://www.ebi.ac.uk/ena/browser/view/PRJEB36951. Genome assemblies for additional *E. coli* strains used here are available from NCBI: ([MG1655](https://www.ncbi.nlm.nih.gov/nuccore/U00096), [SE11](https://www.ncbi.nlm.nih.gov/nuccore/AP009240.1), [REL606](https://www.ncbi.nlm.nih.gov/nuccore/CP000819.1), [CFT073](https://www.ncbi.nlm.nih.gov/nuccore/AE014075.1), [W](https://www.ncbi.nlm.nih.gov/nuccore/CP002967.1), [IA136](https://www.ncbi.nlm.nih.gov/nuccore/CU928164.2), [O157:H7-EDL933](https://www.ncbi.nlm.nih.gov/nuccore/AE005174.2))

## Introduction

Bacterial genomes are extremely dynamic, undergoing frequent loss and gain of both chromosomal and extrachromosomal elements including insertion sequences, rep sequences, ICE elements, and plasmids (Touchon et al. 2009; Lee et al. 2016; Baltrus 2013; Horesh et al. 2021). Quantifying these rapid genome dynamics is critical for understanding population genetic processes, such as the generation of genetic variation and the action of selection. However, it is difficult to quantify such dynamics without accurate and complete (fully contiguous) genome assemblies. Unfortunately, the most dynamic genome elements tend to be both repetitive and plentiful in bacterial chromosomes and plasmids, and thus the most likely to lead to genome mis-assemblies.

Until recently, the most common method for assembling bacterial genomes was using short reads only (e.g. Illumina). However, it is usually not possible to achieve completely contiguous genome assemblies using short reads alone. The development of long read technologies (PacBio and Oxford Nanopore) now allows routine fully contiguous assembly of bacterial genomes (Koren and Phillippy 2015; Wick et al. 2017b). One shortcoming of long read technologies is that they can have low accuracy (although this is rapidly changing). Because of this, one of the most common means of building high-accuracy reference-level genomes is to use hybrid assembly methods, which combine the accuracy and economy of short read technology (e.g. Illumina) with long reads from the economical and accessible Oxford Nanopore (ONT) platform.

Considerable work has been done to address the problem of bacterial genome assembly, using either long-read only or hybrid approaches. Wick and Holt have performed extensive benchmarking of long-read assemblers using both simulated and real ONT read datasets (Wick and Holt 2019), finding that the long read assemblers Flye and Raven perform well across a number of datasets. However, the number of different genomes used in that study (six) was not extensive, and thus may not tell the full story of the strengths and weaknesses of specific assemblers. De Maio et al. (De Maio et al. 2019) evaluated assemblies for 20 Enterobacteriaceae genomes, including four *Escherichia coli* (*E. coli*) using hybrid data, either Illumina and PacBio, or Illumina and ONT. However, the only long-read assembler assessed was Flye.

Here we use ONT and Illumina data from 49 phylogenetically diverse *E. coli* strains, with genome sizes ranging from 4.5 Mbp to 5.4 Mbp, to assess differences in hybrid assembly accuracy for five assembly methods (with polishing when necessary): Unicycler (Wick et al. 2017a), Raven (Vaser and Šikić 2021), Redbean (Ruan and Li 2019), Canu (Koren et al. 2017), and Flye (Kolmogorov et al. 2019). Unicycler is unique in this group in that it first constructs a short read assembly graph, and then resolves ambiguities in the graph using long reads; the latter four are among the most accurate long read assemblers (Wick and Holt 2019).

To quantify assembly accuracy in the absence of known “ground truth” assemblies, we use five metrics designed to measure both structural accuracy and sequence accuracy. We assess structural accuracy by quantifying (1) the contiguity of chromosomes and plasmids; (2) the fraction of concordantly mapped Illumina reads that have been withheld during assembly; and (3) whether all rRNA operons are correctly oriented. We assess indel and substitution frequency by quantifying (1) the fraction of open reading frames that appear truncated and (2) the number of variants that are called using Illumina reads only.

Our results highlight the fact that some assemblers perform predictably better than others. In addition we find that significant differences in assembly accuracy arise because of unidentified strain-specific characteristics that affect long read consensus accuracy. This suggests that currently, there are fundamental limitations to assembly accuracy for bacterial genomes when relying on ONT and Illumina data. At the same time, they emphasise the utility of consensus assembly approaches such as those implemented by Tricycler (Wick et al. 2021).

## Results and Discussion

In order to test the utility of the proposed assembly metrics, we calibrated the metrics on a known “ground truth” *E. coli* K12 MG1655 reference genome. We first obtained ONT and Illumina data for our laboratory strain of MG1655. To ensure that our lab strain matched the sequence of the NCBI reference MG1655 genome (and could truly serve as ground truth), we first called variants against the NCBI reference sequence using Illumina reads alone and the breseq pipeline, which identifies variants with strong statistical support (Deatherage and Barrick 2014). breseq identified ten differences (including substitutions, indels, and structural changes) between our lab strain and the reference MG1655. Seven of the ten changes were shared with ATCC strain 700926 (**Table S1**), which is provided by ATCC as having the MG1655 genotype, but is known to differ from the NCBI reference sequence (Freddolino, Amini, and Tavazoie 2012). This suggested that our lab strain is derived from ATCC 700926.

We incorporated these ten changes into the reference genome sequence, and designated this as ground truth. We note that given the relatively conservative nature of breseq, there may be SNPs present in our lab strain that were not identified (false negatives), although it is unlikely that there are a large number of these.

We next used the MG1655 ONT and Illumina data to produce five hybrid assemblies. These assemblies were constructed with the same Illumina data used for the breseq analysis above, as well as an additional 500 Mb of ONT data (approximately 100x coverage). We then compared each of these assemblies to our ground truth MG1655 sequence. For all assemblers, we found a single contig, all rRNA operons correctly oriented, and the vast majority of paired-end Illumina reads concordantly mapped (99.4% for all). Thus, using these metrics, we would infer that the correct genome structure had been obtained by all assemblers. However, direct comparison with the reference showed that the Canu assembly had duplicated sequences at both the start (34.3 Kbp) and end (63.4 Kbp) of its assembled contig, and the Redbean assembly had 3.7 Kbp gap at the end of its contig, illustrating that these metrics do not detect all changes to genome structure, especially when present at the ends of contigs.

We also inferred SNP and indel frequency in the MG1655 assemblies. We first quantified the fraction of truncated ORFs, which should correlate with the number of indels present in the assembly (see **Methods**). We found between 142 (3.3%, Unicycler) and 268 truncated ORFs (6.1%, Flye) in each assembly (**Table 1**). A small number of truncated ORFs is expected, as pseudogenes are common in *E. coli*; performing the same analysis on the MG1655 reference identified 142 out of 4318 ORFs as truncated (3.3%). Finally, we used the breseq pipeline to call variants in the assemblies using the Illumina reads only. We found between four (Unicycler) and 128 (Flye) SNPs and indels, with the vast majority of errors being deletions. These results suggested that the Unicycler assembly was more than an order of magnitude more accurate in terms of sequence accuracy than the other assemblies.

**Table 1.**
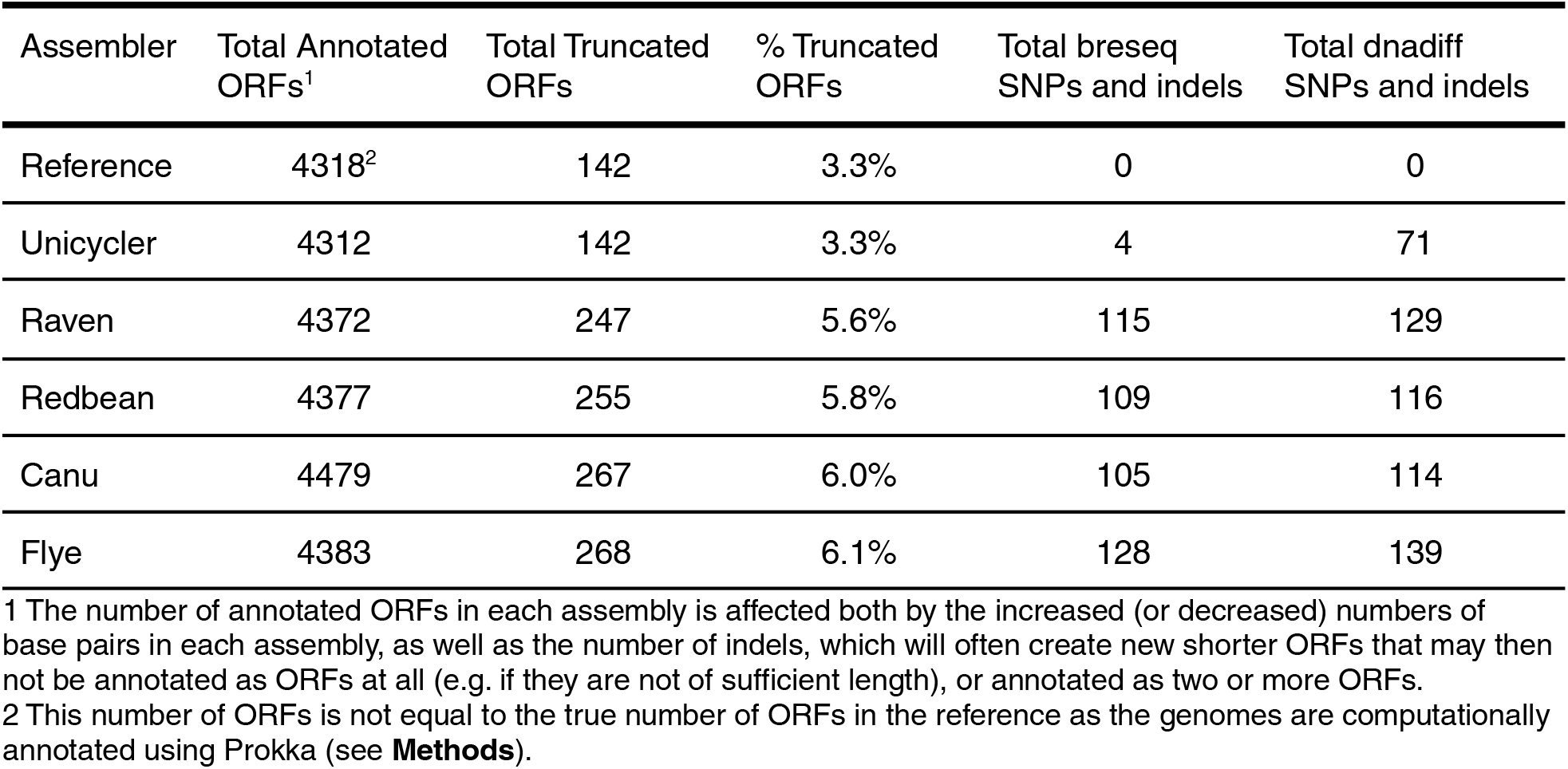
Percentage of truncated ORFs and total SNVs in different MG1655 assemblies relative to ground truth.

However, the breseq pipeline conservatively calls SNPs and indels, so we also compared the assemblies to the original ground truth sequence using dnadiff (Kurtz et al. 2004), which performs a full genome alignment to determine sequence differences. This analysis revealed that all five assemblies contained more SNPs and indels than we discovered using breseq. The most accurate assembly, Unicycler, contained 9 indels and 62 substitutions relative to the reference. Notably, 69 of the 71 differences were in rRNA operons, and are likely due to the repetitive nature of these regions (there are seven rRNA operons in *E. coli*), and difficulties in accurate polishing. However, this is a difficulty that may be resolved by more carefully considered polishing steps (Wick and Holt 2021).

Using dnadiff to compare the other assemblies to the ground truth reference, we found they contained between 114 (Canu) and 139 (Flye) SNPs and indels. These results highlight the fact that even for relatively small bacterial genomes, obtaining highly accurate and complete hybrid genome assemblies is not easily done. In addition, they show that one advantage of long read-first assemblers compared to short read-first assembly (i.e. Unicycler) is that there is a considerably lower rate of error within repetitive regions. This is starkly indicated by the fact that the Unicycler assembly had approximately 20-fold more errors called by dnadiff (which relies on whole genome alignment SNP and indel calling, and thus includes repetitive regions) compared to breseq (which relies on short read variant calling and largely ignores repetitive regions). In contrast, the number of errors in the long read-first assemblies was similar for both dnadiff and breseq (**Table 1**).

### Genome sequencing and assembly of phylogenetically diverse *E. coli* strains

To more thoroughly investigate the accuracy and consistency of bacterial genome assembly methods, we produced assemblies for a set of 49 phylogenetically diverse *Escherichia coli* strains of varying genome complexity and size (Ishii et al. 2006; Breckell and Silander 2020) (**Fig. 1**). We obtained at least 30-fold coverage of 250bp paired end Illumina data for 47 of the 49 strains, and more than 30-fold coverage of 100 bp paired end data for the remaining two. We also obtained a median of 770 Mbp of Oxford Nanopore (ONT) data for each strain (IQR: 545 Mbp - 1118 Mbp). The median average read length across all strains was 5.3 Kbp (IQR: 3.9Kbp - 6.2 Kbp), and median quality scores for all reads across all strains was 13.3 (IQR: 12 - 14). We filtered the ONT reads using Filtlong to retain 500 Mbp (approximately 100-fold coverage), prioritising quality over length (**Methods**). The filtered data sets had a median average read length of 7.7 Kbp (IQR: 6.3 Kbp - 8.0 Kbp; **Fig.2A**) and a median quality score of 14.6 (IQR: 14 - 15; **Fig. 2B**). Surprisingly, we found that both read length and quality varied considerably between strains, despite the data being produced by identical genomic DNA prep methods, flow cell chemistry, and basecalling software.

**Figure 1.**
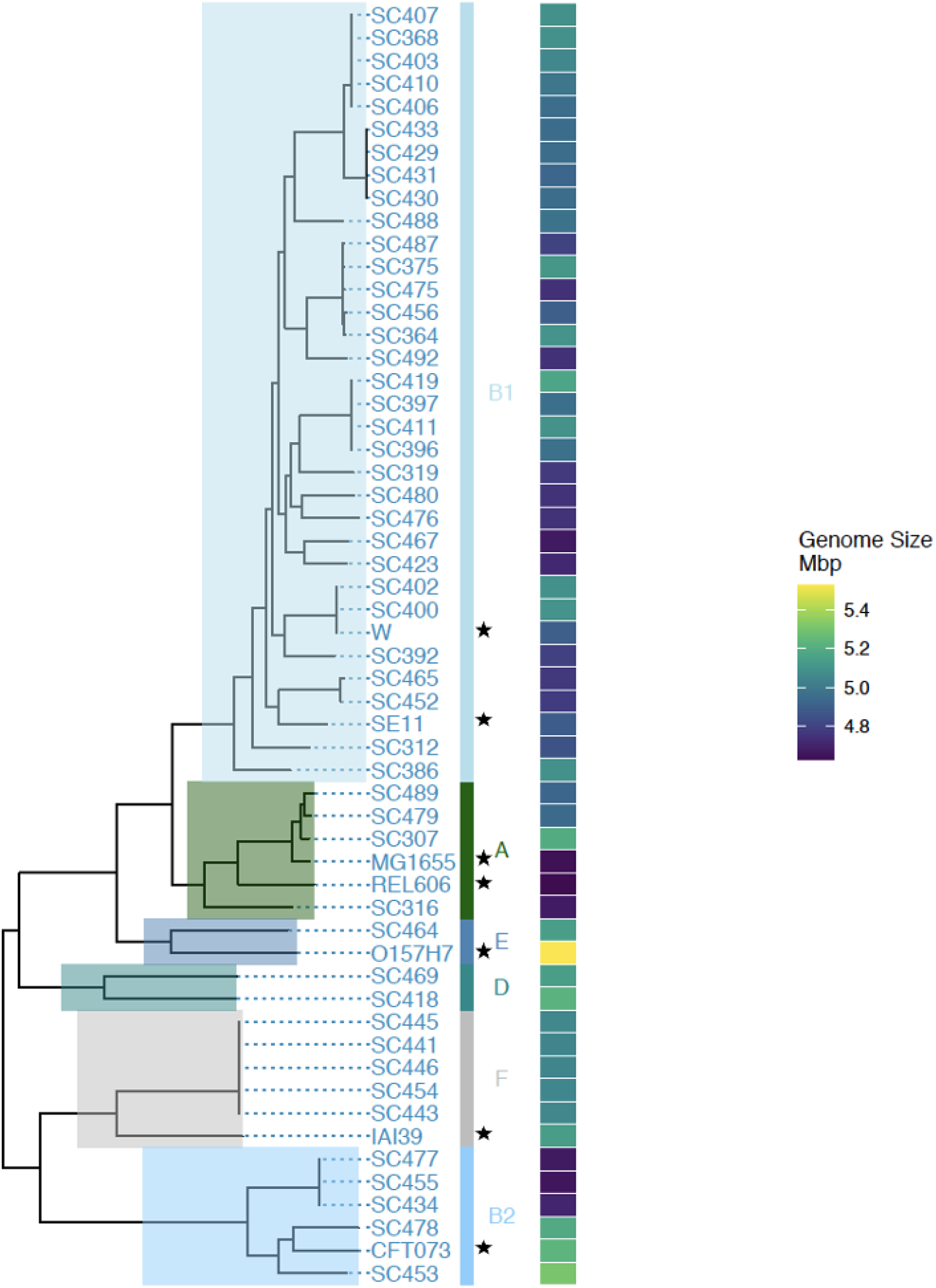
Phylogeny of the *E. coli* strains used to assess assembly accuracy. We used 49 strains to test assembly methods, with representatives from all the major clades of *E. coli* (A, B1, B2, D, E, F). For guidance, we have included seven well known *E. coli* strains in the phylogeny (MG1655, REL606, W, SE11, O157:H7, IAI39, and CFT073), indicated by stars. The 49 strains we use have a wide variety of genome sizes, ranging from 4.5 Mbp to 5.4 Mbp, indicated in the heatmap on the right side of the phylogeny (genome sizes are the means of all five assemblers and include all contigs).

**Figure 2.**
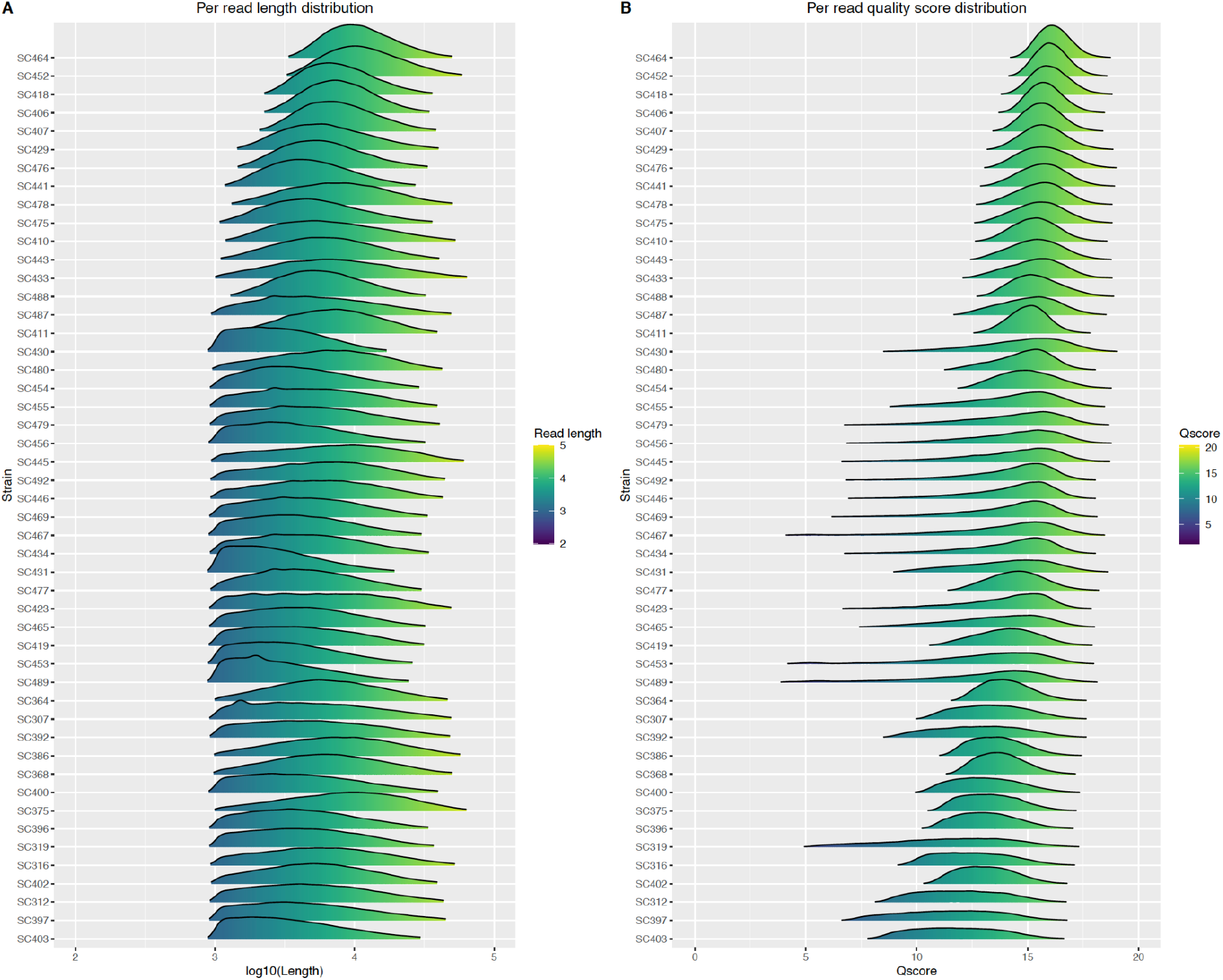
ONT read length and read quality score distributions for all filtered ONT datasets, ordered by percentage of reads with a quality score greater than 15. ONT reads were filtered prior to assembly to retain approximatel 500 Mbp using filtlong, prioritising read quality over length. **A. Filtered read length distributions for each strain. B. Filtered per read average quality scores for each strain**.

We used this data to produce hybrid assemblies with the five assemblers: Canu, Flye, Raven, Redbean and Unicycler. As with the MG1655 assembly, we polished each long read-only assembly (Canu, Flye, Raven, and Redbean) using four rounds of Racon with ONT data, followed by Pilon and Racon with the Illumina reads (see **Methods**). It has been previously reported that Canu assemblies do not necessarily benefit from polishing with long read data (Goldstein et al. 2019). However, we polished all long read assemblies in the same way for consistency. As Unicycler contains a built-in polishing step, we did not polish these assemblies.

### Structural Accuracy and Consistency

We first quantified the total number of contigs in each assembly. Across all assemblers and strains, a median of two contigs were produced (IQR 1-4). Raven consistently produced the fewest contigs, followed closely by Flye. Unicycler, Canu, and Redbean assemblies were the most fragmented (**Fig. 3A** and **Supp. Fig. 1**).

**Figure 3.**
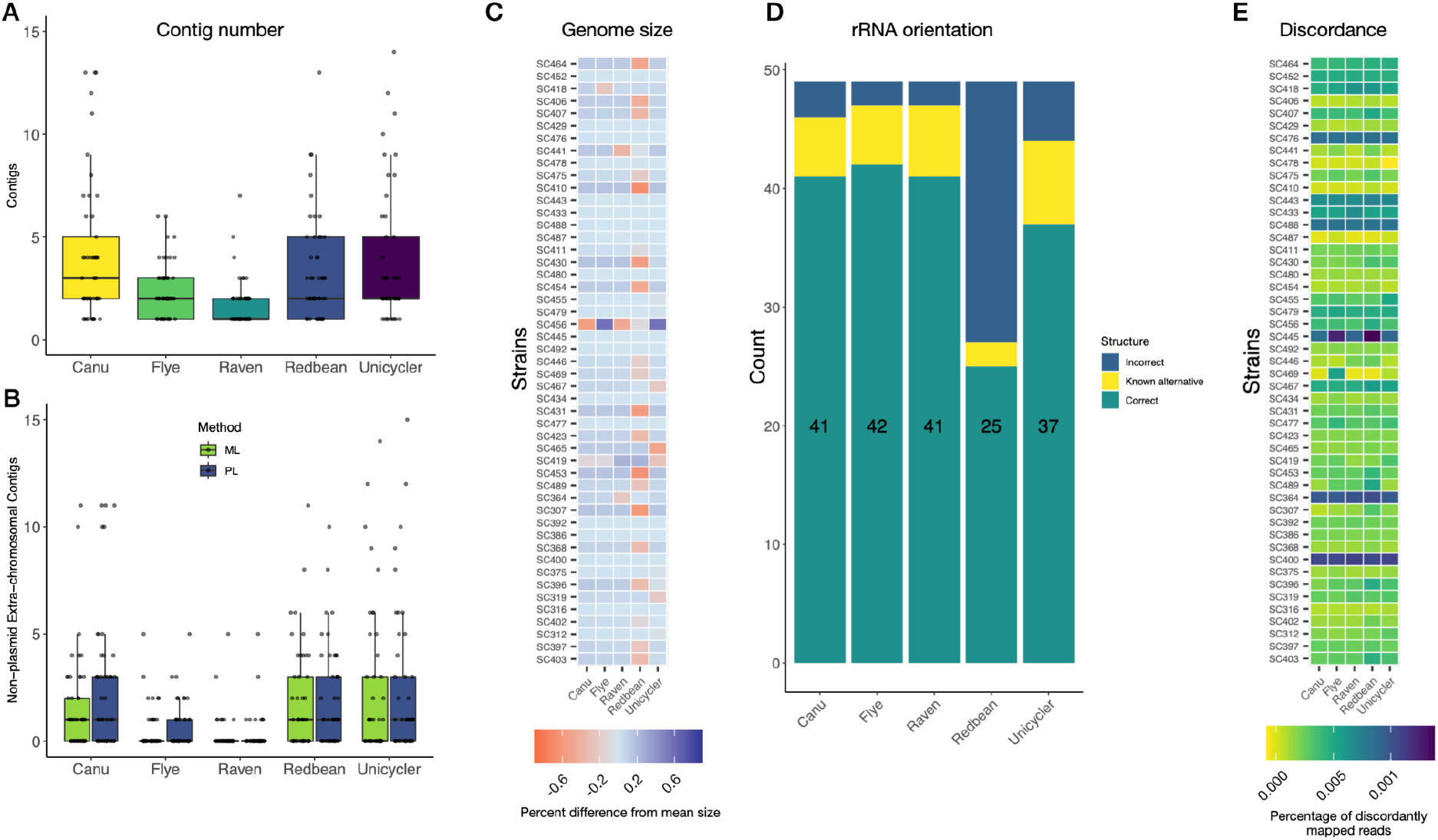
Genome structural contiguity, consistency, and accuracy across assemblers. **A. Total number of contigs for each strain and assembler**. Most strains assembled into five or fewer contigs, with raven consistently having the fewest. **B. Total number of non-plasmid contigs for all strains across assemblers.** We identified plasmid contigs using MLplasmids and Plasmid finder. As in panel **A**, raven assemblies consistently produced the fewest non-plasmid contigs, with the vast majority of assemblies consisting of a single chromosomal contig. **C. Difference in chromosomal assembly size for each assembler.** The difference in assembly size for each assembler from the mean of all assemblers is indicated. Differences in assembly length were generally less than 0.1% of the total genome size, equivalent to 5,000 bp for a 5 Mbp chromosome. Redbean assemblies were often the shortest, in many cases by approximately 0.5% (25 Kbp for a 5 Mbp genome). Strains are ranked from top to bottom by the percentage of reads with a quality score greater than 15 (see **Fig. 1**). **D. rRNA operon orientation for all strains across assemblers.** We tested whether each of the seven rRNA operons were in an orientation that has been observed previously in E. coli. Flye, Raven, and Canu consistently produced structurally correct assemblies with expected or known alternative rRNA operon orientations (47 out of 49 genomes in the case of Raven and Flye). In contrast, Redbean produced structurally correct assemblies for only 27 out of 49 strains. **E. Illumina mapping discordancy for each assembler.** We withheld 5,000 pairs of Illumina reads from each assembly, mapped these back on to the assembled genomes, and calculated the fraction of mapped reads that were discordantly oriented. We observed strain specific trends in discordant mapping, but no assembler-specific trends.

These results suggested that Raven and Flye were optimal for maximising assembly contiguity. However, it is also possible that the additional contigs in the more fragmented assemblies were true (extrachromosomal) plasmid contigs, which for Raven and Flye had been incorrectly incorporated into the chromosome. We thus classified each contig as plasmid or chromosomal using mlplasmids (Arredondo-Alonso et al. 2018) and PlasmidFinder (Carattoli et al. 2014). These methods can identify small contigs as plasmids; however, they cannot identify regions of chromosomal contigs as plasmid. Both classification methods identified many of the small contigs in the Canu, Unicycler, and Redbean assemblies as chromosomal. In contrast, almost all small contigs in Flye and Raven were identified as plasmids **(Fig. 3B)**. This suggests that the additional contigs are in fact unincorporated chromosomal fragments.

We next examined consistency in the size of each genome assembly across all assemblers. We calculated the mean size of the largest contig for each strain for all assemblers (under the assumption that this was the chromosomal contig) and tested whether there were systematic deviations from this mean size for each assembler. Although this is a relative metric and does not objectively establish whether one assembler is the most accurate, it provides insight into whether specific assemblers tend to arrive at the same or different results - a “wisdom of the crowds” approach. We found that the median difference in assembly size across all strains and assemblers was 39 kbp (0.0078%). The most consistent pattern was that shorter assemblies were frequently produced by Redbean (**Fig. 3C**). This is expected, as Redbean generally produced more fragmented assemblies that had parts of the chromosome as unincorporated contigs. Although previous work has suggested that Canu assemblies are often larger as they often contain overlaps at the start and end of chromosomes (Wick et al. 2017a), we did not observe this as a general pattern.

Single-contig circularised genomes may be structurally inaccurate, containing large inversions, deletions, insertions, or amplifications. One means of identifying structural mis-assemblies is through examining the order and orientation of rRNA operons. *E. coli* contains seven highly conserved rRNA operons. If mis-assembly occurs at these locations, these operons may be joined incorrectly to the neighboring chromosomal regions (in orientation, chromosomal location, or both). We used Socru (Page, Ainsworth, and Langridge 2020) to assess whether rRNA operons were properly oriented within each assembly. We found that Flye produced the greatest fraction of assemblies having all seven rRNA operons correctly oriented or in a known alternative orientations (42 out of 49 in the expected orientation and five out of 49 in a known alternative orientation). Raven and Canu were similar, with 47 and 46 rRNA operons, respectively, in the expected or known alternative orientations (**Fig. 3D**); Unicycler had slightly fewer. However, for the Redbean assemblies, only 27 out of 49 had all operons in an expected or known alternative orientation.

We also assessed structural accuracy by mapping 5,000 paired end Illumina reads (10,000 individual reads) that we withheld during assembly (see **Methods**). We then quantified the fraction of discordantly mapped reads (i.e. the fraction of all mapped paired-end reads that were not concordantly oriented). This should yield insight on whether there are frequent small scale inversions or duplications in the assemblies. We found that across all assemblers, very few mapped discordantly, with a median of 0.26% (approximately 26 reads; IQR 0.14% to 0.36%; **Fig. 3E**). Although Redbean performed very slightly worse than the other assemblers, almost all differences in discordance were strain-specific rather than assembler-specific, suggesting that some strains are simply harder to assemble correctly, perhaps due to the arrangement or numbers of repetitive elements in the genome. Alternatively, the Illumina sample libraries may have differed in the fraction of artifactual chimeric reads (Marçais, Yorke, and Zimin 2015).

### Sequence Accuracy

Large-scale structural accuracy is only one assembly characteristic; accuracy at the base pair level is also a critical aspect, especially as ONT data is prone to systematic indel errors in homopolymeric regions (usually deletions). These indels can lead to frameshifts, resulting in premature stop codons and shortened open reading frames (ORFs). To estimate the extent of indels in each assembly, we quantified the percentage of unexpectedly short ORFs (see **Methods**). In the MG1655 reference genome we found that 3.3% of all ORFs were truncated (**Table 1**); repeating this analysis for each of the 49 other *E. coli* strains we observed that Unicycler assemblies consistently had the smallest fraction of truncated ORFs (**Fig. 4A**). These results suggest that even if long read-first ONT assemblies are polished with short reads, this does not completely mitigate the problem of frequent indels.

**Figure 4.**
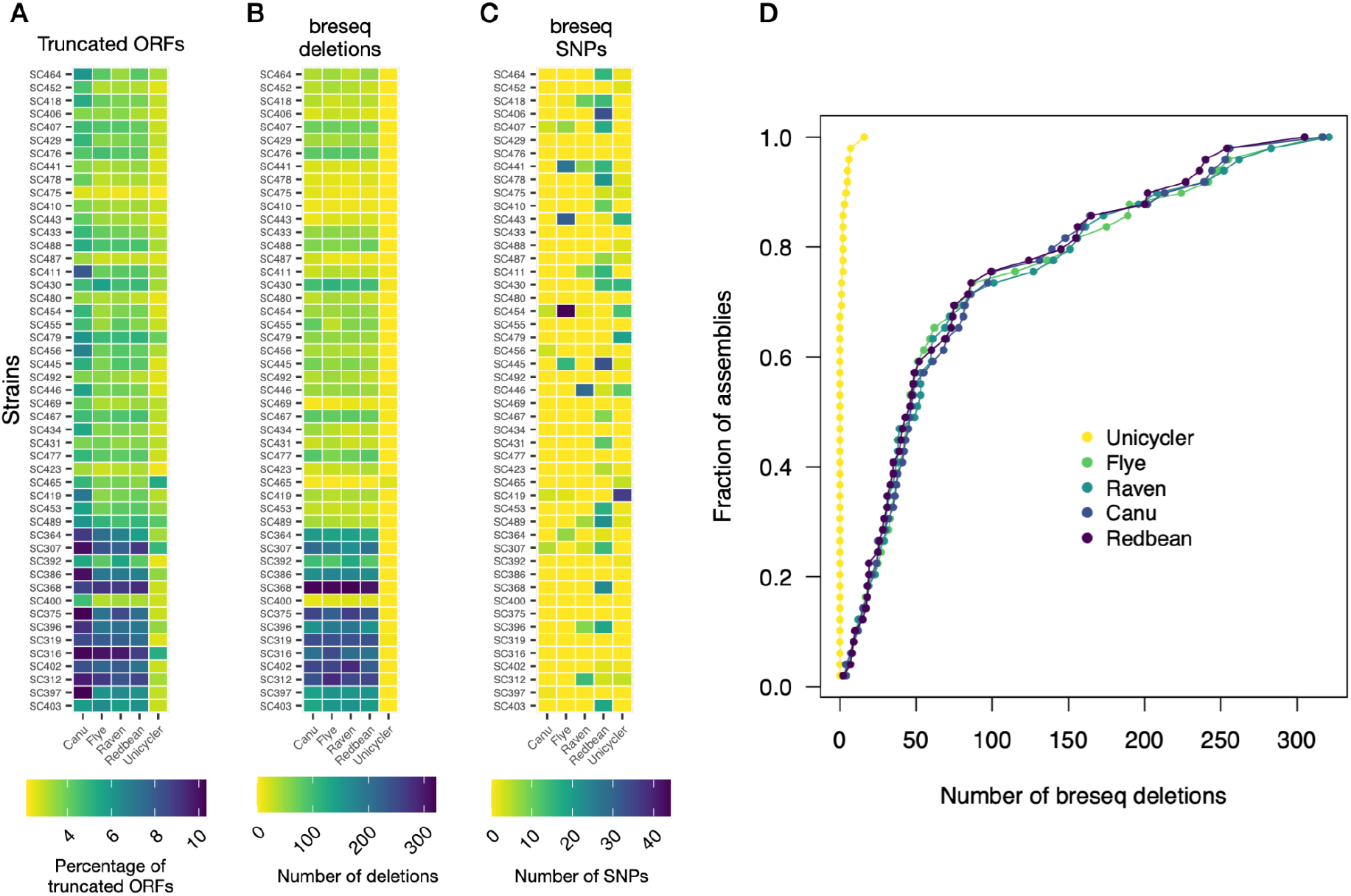
Genome sequence accuracy metrics across assemblers. **A. Percent of truncated ORFs for each assembler.** We calculated the fraction of truncated ORFs for each assembler as those ORFs that were 90% or less of the expected length (see **Methods**). We found that Unicycler consistently exhibited a far lower fraction of truncated ORFs. **B. Number of deletions in each assembly called via breseq using Illumina reads alone.** We used the breseq pipeline to call deletions using Illumina reads. Unicycler assemblies consistently had the fewest deletions; this is somewhat expected as the initial Unicycler assembly was constructed using Illumina reads only. However, all other assemblies were also polished using the same set of Illumina reads. **C. Number of SNPs in each assembly called using Illumina data alone.** We found few SNPs using breseq for any assembler, although in a small number of cases, we observed up to 40 SNPs. These tended to be both strain and assembler-specific, rather than consistent for any one assembler or any one strain. **D. Cumulative plot of the number of breseq-called deletions for each assembler.** The majority of long read-first assemblies have fewer than 50 deletions, far more than Unicycler. However results from ground-truth MG1655 comparison suggest that there are few false negative errors in long read-first assemblies, in contrast to Unicycler.

While the fraction of short ORFs gives some insight into the prevalence of deletions, there are also real differences between strains in the number of pseudogenes, which affects the fraction of short ORFs. In an effort to reduce the effects of these differences on our assessment of assembly accuracy, we used the breseq pipeline to call SNPs and indels in each assembly using Illumina reads only. In all cases, we used the same Illumina reads that were used in the original assembly or subsequent polishing steps. We found that while on average few indels or SNPs were identified (median of 37 deletions, IQR 9 - 81; median of 0 SNPs, mean 2.9), some assemblies had considerably more - between 200 and 300 deletions and up to 40 SNPs **(Figs. 4B-4D)**. Again we found that Unicycler assemblies contained the fewest deletions across all strains. This is perhaps not surprising, as the initial Unicycler assemblies were made using these same reads. It is critical to note though that both the truncated ORF metric and the breseq variant calling metric may be prone to missing real SNPs and indels present in Unicycler assemblies, as they tend to occur in repetitive regions such as rRNA, as observed above for MG1655,. These errors do not necessarily affect ORF length, nor are they called by breseq. However, it is difficult to identify these false negatives with certainty in the absence of a ground truth genome. This contrasts with the long read-first assemblies, which do not appear as prone to false negative errors when assessed for quality using breseq variant calls.

Nevertheless, we sought to test the possibility that false negative indels and SNPs were present in the Unicycler assemblies by comparing them to the other long read-first assemblies. We hypothesised that if indels and SNPs in Unicycler assemblies were present primarily in repetitive regions, they should be clumped along the genome in these regions compared to other assemblers. We tested this by taking the Raven assembly for each strain as a “reference,” and aligning all other assemblies against it in a pairwise fashion. We selected the Raven assemblies as the reference because they were consistently among the most accurate. From these alignment, we could infer mismatches in the assemblies, which likely arise from errors in either the Raven assembly, or in the assembly aligned to Raven. We found four different patterns in the genome-wide error profiles. For some strains, the Unicycler assemblies consistently exhibited fewer SNPs and indels across the genome (**Fig 5A**). In these cases, the slope of the cumulative number of SNPs and indels in the Unicycler assemblies was approximately half the slope of the other assemblers, which is expected if the Unicycler assembly generally contains few errors - the cumulative difference in SNPs and indels between the Unicycler and Raven assemblies are errors in the Raven assembly. The slope of the cumulative error curves for all other assemblies is approximately twice the Unicycler slope because they have similar numbers of errors as the Raven assemblies spread across the genome, so the cumulative error curves reflect errors in both the Raven and the other assemblies.

**Figure 5.**
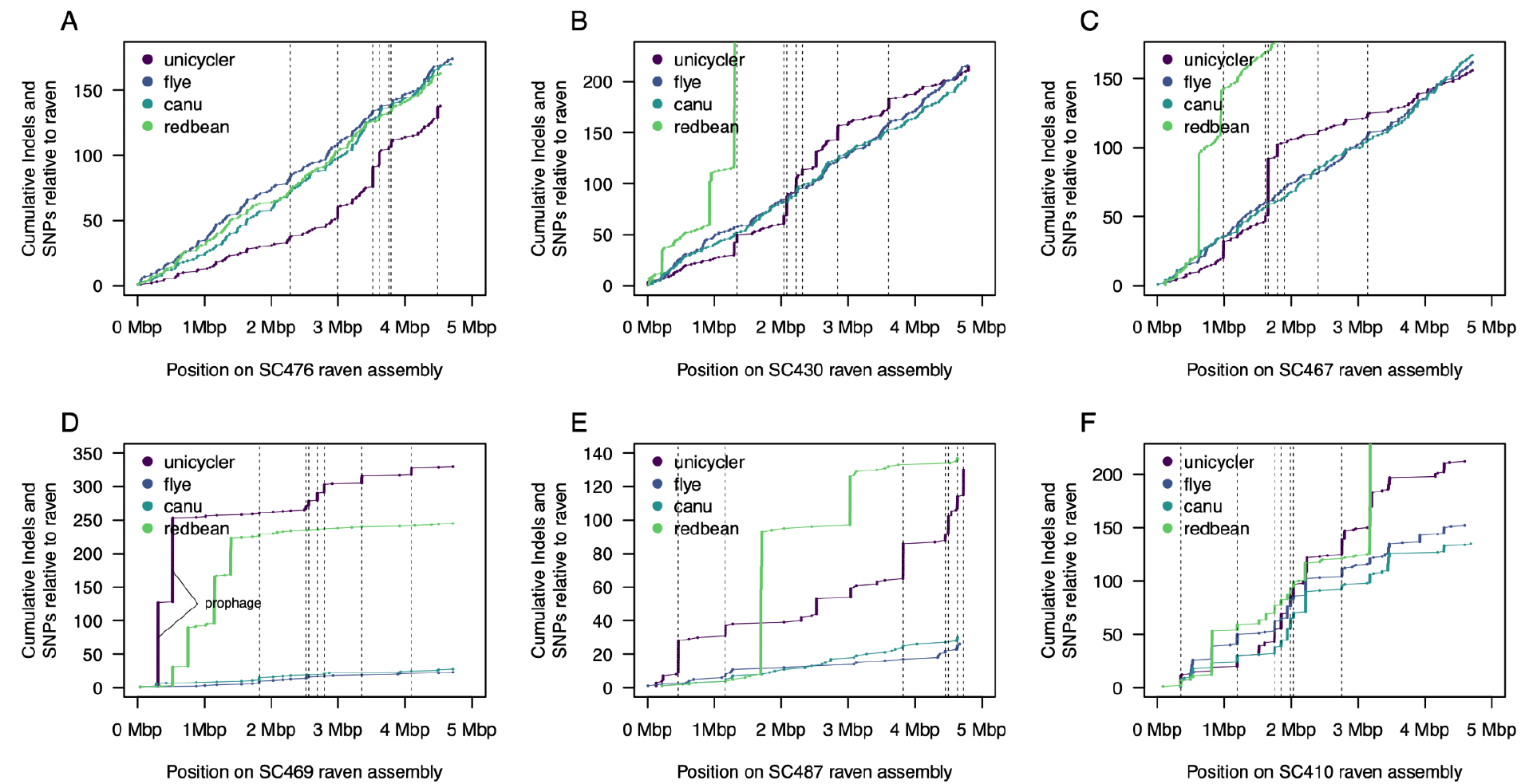
Whole genome alignments for all assemblies for six example strains. For each strain, we used the Raven assembly as a reference, aligned all other assemblies against it, and identified SNPs and indels using dnadiff. Each panel shows the cumulative number of SNPs and indels for each assembler along the genome of one strain. The location of the rRNA operons are shown as dotted vertical lines.The locations of two prophages are indicated in (**D**), where the Unicycler assembly exhibits a large number of errors. In (**B**), (**C**), and (**F**), the Redbean assemblies contained a far larger number of SNPs and indels than the other assemblies, so the complete cumulative curves are not shown.

The second pattern we observed was one in which similar total numbers of SNPs and indels appeared for all assemblers, but in Unicycler (and often Redbean), the majority were due to concentrated stretches of errors (**Figs. 5B** and **C**). In these cases, the large numbers of consecutive errors in the Unicycler assemblies tended to occur at rRNA regions, confirming that when errors occur within Unicycler assemblies, they tend to accumulate in repetitive regions..

The third pattern we observed was one in which most long read assemblers exhibited very few differences, but Unicycler and Redbean exhibited long stretches of errors, again in rRNA regions or other repetitive elements (**Figs. 5D** and **E**). Finally, in some cases we found that errors tended to be clumped for all assemblers, but distributed across the genome (**Fig. 5F**). When these errors are present in the “reference” Raven genome, these are apparent as a concerted step increase in the cumulative number of errors for all assemblers; when these are present in only one assembler, this is apparent as a step increase only for that assembler. These results show conclusively that although at first glance the Unicycler assemblies are highly accurate in terms of SNPs and indels, there are a large number of errors that are present in repetitive regions but which are not easily identified found by quantifying the fraction of truncated ORFs or by calling variants using breseq.

### Effects of data quality on assembly accuracy

In addition to assembler-specific errors, we expected that differences in assembly accuracy might be dependent on the quality of the input data, which often differed between strains. Indeed, we observed that differences in accuracy were often strain-specific rather than assembler-specific. For example, some strains exhibited consistently higher discordance (**Fig. 3E**) or systematically higher indel numbers (**Fig. 4B**). These differences have two possible sources - the quality of input data, or characteristics intrinsic to each strain that preclude accurate assembly. Both the Illumina and ONT data quality differed between strains in read length and quality (**Fig. 2** and **Supp. Fig. 2**). Variation in ONT read length was almost certainly due to subtle differences in sample treatment during genomic DNA prep. However, we did not expect substantial variation in ONT read quality, as library chemistry, flow cell chemistry, and basecalling methods were identical for all strains (9.4.1 flow cells with RBK004 chemistry and guppy 4.2.2). Surprisingly, we observed differences in read quality even for library samples run on the same flow cell and on the same day.

We first tested for effects of input data quality on assembly structural accuracy by quantifying correlations between ONT read length (read N50) and any accuracy metric. We found no strong correlation between read length and any of the structural quality metrics **(Supp. Fig. 3A-D)** for any assembler, although Redbean assemblies showed a trend for lower structural accuracy as assessed by rRNA operon orientation (**Supp. Fig. 3A**, rightmost panel). We also found no correlation between read length and any sequence accuracy metric (**Supp. Fig. 4**).

We also tested for correlations between ONT read quality (**Fig. 2B**) and assembly sequence accuracy. The result from the initial set of assemblies suggested that strains with lower read qualities exhibited both increased levels of truncated reading frames and breseq-called indels (**Fig. 4**). However, this pattern was not clearcut. For example, strain SC400 exhibited very few deletions or truncated reading frames despite having relatively low quality ONT data. Interestingly, this strain is very similar to strain SC402 both phylogenetically and in genome size (**Fig. 1**), but strain SC402 had considerably more truncated ORFs and indels (fourth row from the bottom in the **Fig. 4** heatmaps) despite having very similar read quality scores. We currently do not have an explanation for this discrepancy.

To clarify the effects of ONT read quality on assembly accuracy, for six strains we collected between three and six additional sets of read data, either using different flow cells, or as independent multiplexed libraries on the same flow cells. These read sets exhibited varying levels of per read quality distributions (**Fig. 6A**). For each of these read sets we then performed assemblies using all five assembly methods, and calculated the sequence accuracy metrics for each (**Fig. 6B - D**). We observed consistent differences in assembly quality between strains, with some strains exhibiting more than 150 deletions and more than 6% truncated ORFs regardless of the ONT read set or assembly method (with the exception of Unicycler assemblies). In other cases, strains with read sets of nearly the same quality exhibited fewer than 30 deletions on average and less than 4% truncated ORFs. This strongly suggests that there are strain-specific characteristics that affect assembly accuracy. As for read length, we also checked for correlations between read quality and assembly sequence accuracy, and found no strong relationship between any metric for any assembler (**Supp. Figs. 5** and **6**).

**Figure 6.**
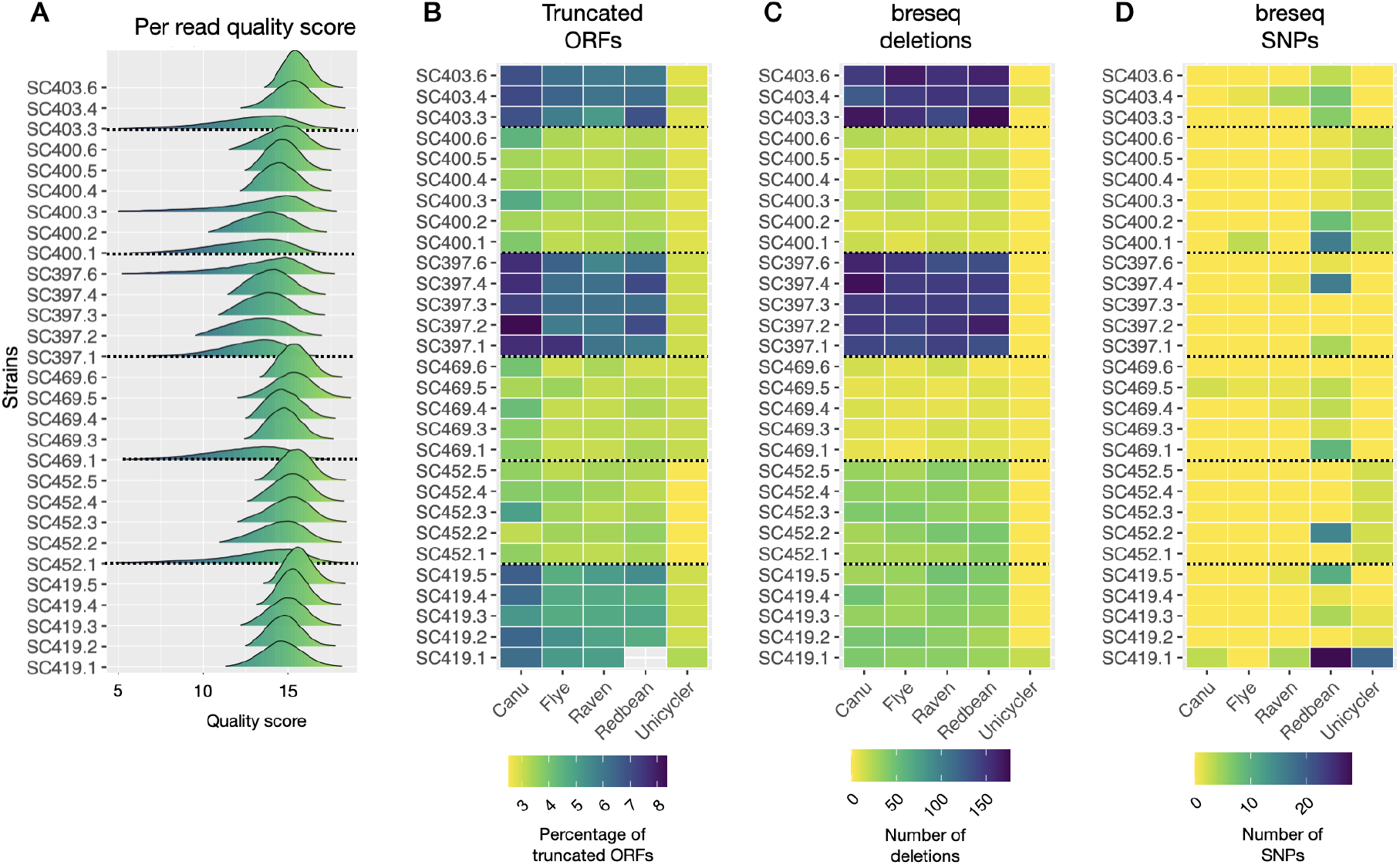
Assembly accuracy is strain dependent and does not correlate with the quality of ONT data. We collected between three and six additional ONT datasets for six strains and then performed assemblies using all five assemblers. **A.** Mean per read quality scores for each data set. The data are arranged by strain, and within strains, by the fraction of reads that are above Q15. **B.** Fraction of truncated ORFs per assembly. **C.** Number of deletions called by breseq. **D.** Number of SNPs called by breseq.

However, all of the assemblers use short read Illumina data at some stage, either for initial assembly (Unicycler) or for polishing, and in the above results, the same Illumina data were used for polishing all of the assemblies for each strain. We thus considered whether the quality of Illumina reads affected assembly sequence accuracy, as there were clear differences in read quality (**Supp. Fig. 2**). Two circumstantial pieces of evidence suggest that Illumina read quality did not have strong effects on assembly sequence accuracy. First, the two strains with the lowest average Illumina read quality, SC455 and SC400, did not exhibit substantially lower assembly metrics (in fact SC400 exhibited better assembly metrics than average). Second, the quality of the Unicycler assemblies, which depend first and foremost on the Illumina reads for assembly, did not show any clear correlation with read quality.

To directly test the effects of Illumina read quality on assembly accuracy, rather than collect additional data, we divided the MG1655 Illumina data into sets of high quality and low quality reads, such that the high quality set had mean quality scores of 38-39, and the low quality set had mean quality scores of 33 to 34 (**Supp. Fig. 7**). We assembled each read set using Unicycler and compared these to the ground truth MG1655 assembly using dnadiff for genome-wide alignment. Surprisingly, we found that the low quality read set resulted in an assembly with the fewest errors (60 SNPs and indels), compared to 65 SNPs and indels for the high quality set and 71 SNPs and indels for the full read set. This suggests that Illumina read quality has little, if any, effect on assembly quality.

Overall, these results suggest that neither Illumina nor ONT read quality (as calculated by the guppy basecaller) strongly affect the resulting hybrid assemblies. Rather, for long read assemblies (but not for short-read first Unicycler assemblies), there are unidentified idiosyncratic characteristics for each strain that affect assembly accuracy, as assessed by quantifying the fraction of truncated ORFs and breseq-called SNPs and indels. There remain two possible explanations for these differences. First, it is possible that some strains have low-accuracy assemblies because they do in fact rely on low quality ONT data, but this quality is not reflected in the quality scores assigned by the base caller. This would manifest as reads having high quality scores but low percent identity to the assembly. Alternatively, some strains may have low-accuracy assemblies because of fundamental problems in creating accurate consensus sequences.

To distinguish between these two possibilities, we examined the relationship between read identity to the assembly and read quality score for three strains that exhibited varying levels of assembly quality (SC403, SC419, and SC452; **Fig. 4**). We found that all three matched almost exactly in the relationship between read identity and quality score, suggesting that the quality score assigned to reads by the guppy basecaller truly reflects the quality of the reads (**Supp. Fig. 8**). These results imply that the primary limitation for accurate assembly is decreased consensus accuracy when doing long read-first assemblies. These intrinsic problems in consensus accuracy are easily diagnosed by comparing assembly metrics between the short-read first Unicycler assembly and the long-read first assemblies. When these metrics are mismatched, this suggests there are problems in consensus read accuracy for the long read-first assemblies.

## Conclusion

Here we have investigated five assembly methods for their ability to generate accurate hybrid genome assemblies from Illumina and Oxford Nanopore data. We have employed three metrics of assembly quality that assess structural accuracy: assembly contiguity, discordancy of short read mapping, and the accuracy of rRNA region arrangement. We have used two metrics to assess sequence identity: the fraction of truncated reading frames observed, the number of indels and SNPs called using short read (Illumina) data.

The data here suggest that Raven and Flye are the top performing assemblers from a structural standpoint, while Unicycler is most effective at minimising indels and substitution errors outside of repetitive regions. These assemblers thus offer complementary advantages. If structural accuracy is critical for the question at hand (for example quantifying genome dynamics for rapidly evolving regions such as IS elements or plasmids), then Raven and Flye are preferable. For other applications, such as evolve and resequence experiments (Long et al. 2015), Unicycler is the preferable method. It is important to note that the metrics used here to assess assembly sequence accuracy likely overestimate the accuracy of Unicycler assemblies - when there are errors in Unicycler assemblies they are most likely in repetitive regions, such as rRNA (**Fig. 5**), and are not easily identified using the assembly accuracy metrics here.

We also found that there are intrinsic properties of certain bacterial strains that limit consensus accuracy. We suggest that by contrasting the qualities of short-read first Unicycler assemblies and long-read first Raven or Flye assemblies, these problems in assembly accuracy can be easily diagnosed. Finally, these results emphasise the importance of consensus assembly methods, such as that offered by trycycler (Wick et al. 2021).

## Methods

### DNA Extraction

We grew single cell colonies from each of the 49 natural isolates overnight in 3 mL of liquid LB media at 37°C. We extracted DNA using either the Promega Wizard DNA extraction kit (following the gram negative bacterial extraction protocol), or phenol chloroform (Quick 2018). We measured DNA size distributions using gel electrophoresis in a 1% agarose gel, and both a Qubit fluorometer and Nanodrop readings to measure the size and quality of each DNA prep.

### Library Preparation and DNA Sequencing

All Illumina Genome sequencing was performed by MicrobesNG (http://www.microbesng.uk) which is supported by the BBSRC (grant number BB/L024209/1).

We prepared ONT sequencing libraries using 400ng of DNA with the SQK-RBK004 or SQK-LSK109 kits according to the manufacturer’s protocol with the following modifications: samples were eluted off Agencourt Ampure XP beads into TE buffer pre-warmed to 50°C; we performed the elution at 50°C; and increased the incubation time in the elution buffer to 10 minutes.

ONT sequencing was carried out on a MinION Mk1b device using R9.4.1 flowcells. 17 flowcells were used in total, with between four and 12 strains run per flowcell (**Supp. Table 2**). Several strains were sequenced multiple times to ensure that at least 500 total Mbp (approximately 100X coverage) with a read N50 greater than 5kb was generated for each strain. For nine strains we achieved complete assemblies with lower coverage (between 24x and 94x), and did not generate additional sequence data for those strains. We defined a complete assembly as having no unidentified contigs, the correct rRNA operon arrangement, greater than 98% illumina read concordancy and fewer than 7% short ORFs.

For each MinION run, we demultiplexed the reads using Deepbinner (Wick 2021a) and basecalled with Guppy v4.2.2.

### Genome Assembly and Polishing

We filtered all Oxford Nanopore reads to retain approximately 500 Mbp using Filtlong (Wick 2021b), specifying the option to use Illumina reads to determine which ONT reads are of high or low quality. From all Illumina read sets, 5,000 pairs of Illumina reads were removed and withheld from assembly (see below).

For all five assemblers (Unicycler, Canu, Flye, Raven, and Redbean) we used default settings. We polished each long read-first assembly with both Illumina and ONT data, using four rounds of Racon with ONT reads, followed by Pilon and Racon with Illumina reads. All basecalling, filtering, assembly and polishing pipelines are available on github (https://github.com/GeorgiaBreckell/assembly_pipeline).

For all long read assemblers we also tested the utility of an intermediate polishing step with medaka v1.2.2 (Oxford Nanopore Technologies 2021). However, we obtained inconsistent results: some assemblies decreased in quality while others changed little. Due to these inconsistencies, we do not report any results with medaka-polishing steps here.

### Multiple Alignment and Phylogenetic Reconstruction

The alignment was created using REALPHY (Bertels et al. 2014) with default parameters and specifying the *E. coli* strains MG1655, REL606, W, SE11, O157:H7, IAI39, and CFT07 as the references. *E. fergusonii* was also used as a reference to allow rooting. RAxML (Stamatakis 2014) was used for phylogenetic reconstruction.

### Quality Assessment

All commands, and software versions used for quality assessment are described in the assembly pipeline available on the github repository linked above. A brief explanation of the tools and their usage is given below.

### Genome Fragmentation

A complete *E. coli* genome should contain a single circular contig representing the chromosome along with a variable number of circularised contigs representing plasmids. We determined fragmentation for each assembly by testing (1) the number of contigs present in each assembly; and (2) whether any additional contigs could be identified as plasmids.

### Plasmid Assignment

We used mlplasmids (Arredondo-Alonso et al. 2018) and Plasmidfinder (Carattoli et al. 2014) to assess whether contigs were of plasmid or chromosomal origin. mlplasmids uses a support vector machine learning approach based on the frequency of 5-mers, with training on taxa-specific chromosomes and plasmids. The result is a posterior probability of a contig belonging to a plasmid or chromosome class. We assigned contigs as plasmids if this posterior probability was greater than 0.5. In contrast, Plasmidfinder uses database matching for contig classification, and returns the identity of the matched plasmid.

### Illumina Mapping Discordancy

We withheld 5,000 Illumina paired end reads from each assembly. We then aligned these to assembly using bwa mem (Li 2013). We used samtools (Li et al. 2009) to extract the fraction of concordantly mapping reads and from this determined the fraction of discordantly mapped reads.

### rRNA Operon Orientation

We used Socru (Page and Langridge 2019) to determine whether the arrangement of inter-rRNA regions in each of our assemblies was as expected, or had been observed previously. Socru maps inter-rRNA regions against its *E. coli* model to determine their presence and arrangement in each assembly. We ran Socru with default parameters and the *Escherichia coli* model of rRNA structure.

### Truncated ORFs

To quantify the number of truncated ORFs we followed the approach outlined in ideel (Watson 2018). Briefly, we annotated each assembly with Prodigal (Hyatt et al. 2010) to produce a set of predicted proteins. We aligned to the Uniprot protein database using Diamond (Buchfink, Reuter, and Drost 2021). We then compared the length of each protein and its top hit. We considered any ORF with a length less than 90% of the top hit as truncated.

### Deletions and SNPs

To quantify the number of SNPs, and deletions in each assembly we used the breseq pipeline with default settings (Deatherage and Barrick 2014). We ran the pipeline with our Illumina reads for each strain, with the assemblies produced by each assembler.

### Pairwise Genome Alignment

We used dnadiff v1.3 (Kurtz et al. 2004) to align whole genomes to quantify differences in SNPs and indels between assemblies of the same strain, using default settings.

## Abbreviations

IQR: interquartile range
ONT: Oxford Nanopore Technologies

## Supplementary Figures and Tables

**Supplementary Figure 1:**
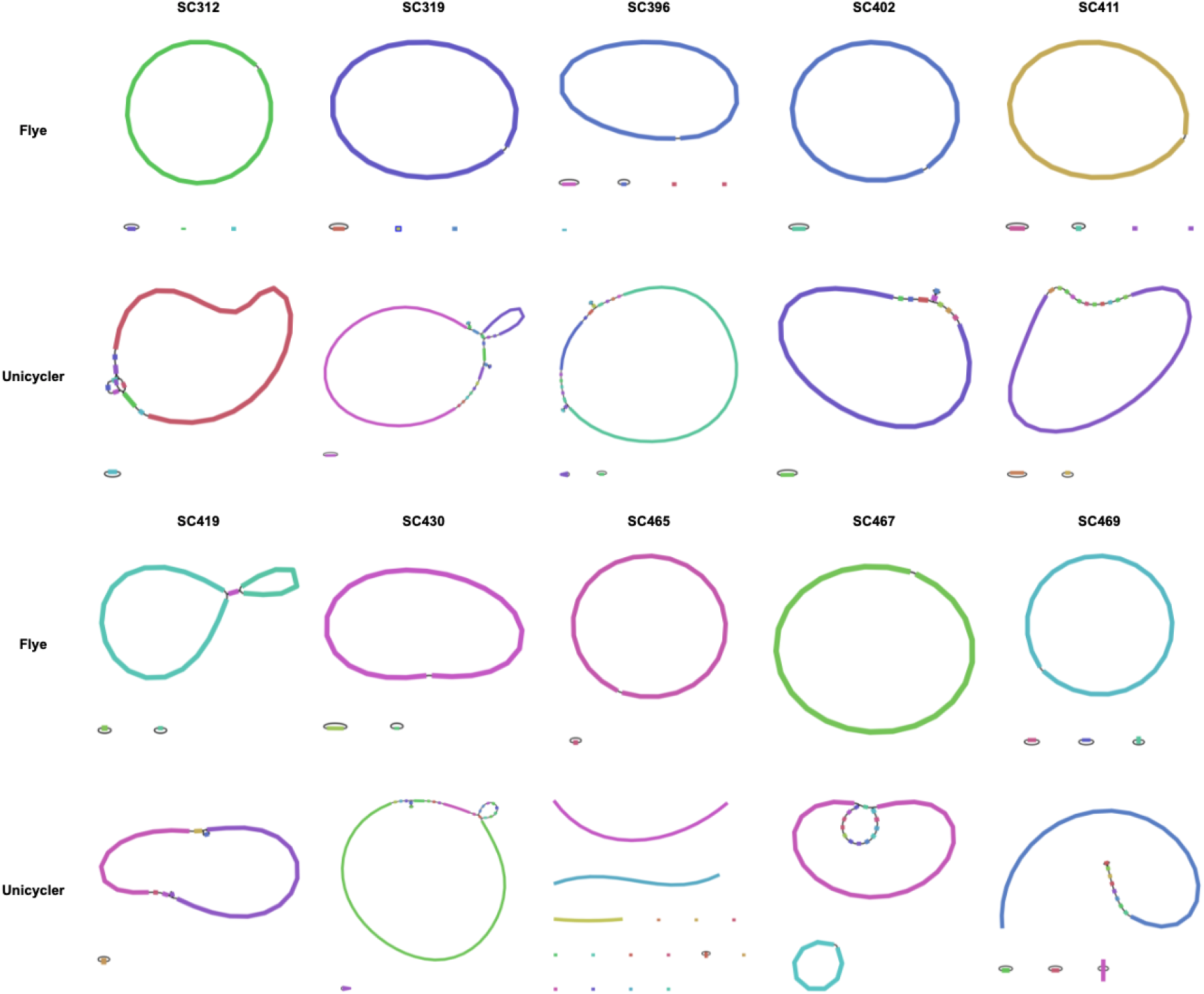
Differences in assembly structure between assemblers. As an illustrative example, here we show bandage (Wick et al. 2015) plots of the 10 most fragmented Unicycler assemblies, coupled with the Flye assemblies for the same strain. Unicycler often produced more fragmented assemblies, while Flye was the second most contiguous assembler. We typically observed the introduction of multiple small contigs within the chromosomal contig of Unicycler assemblies.

**Supplementary Figure 2.**
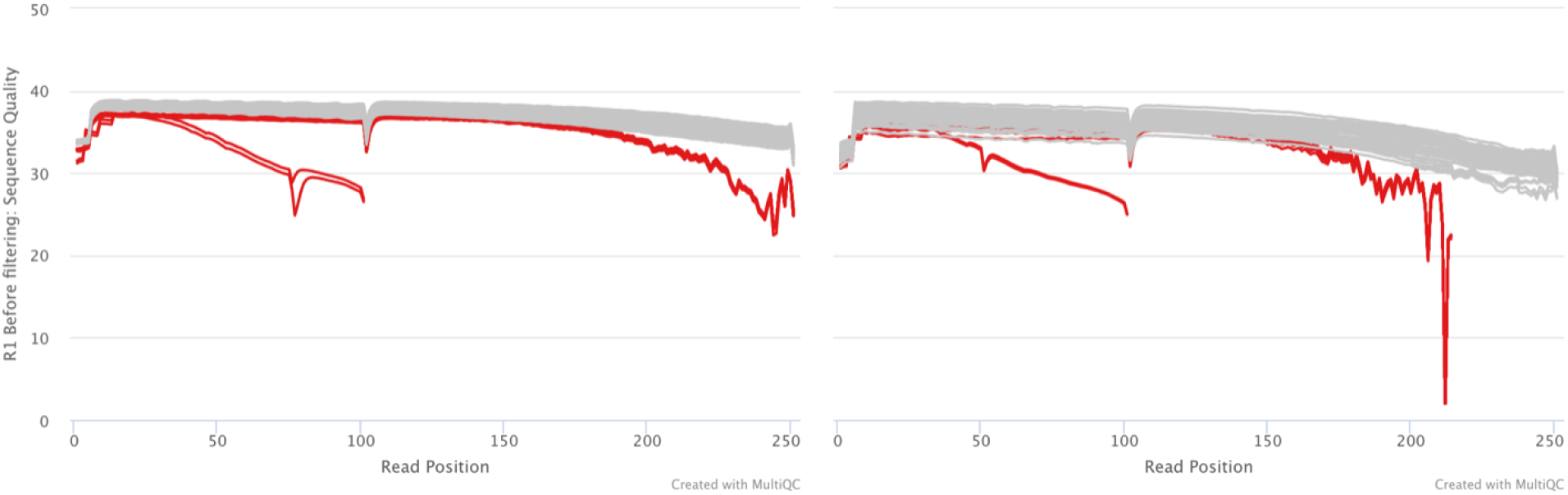
Illumina data was systematically lower quality scores for a subset of strains. Mean quality scores at each position are indicated, calculated using fastp (Chen et al. 2018) and displayed using multiQC (Ewels et al. 2016). Each strain is indicated with a single line, with Read 1 shown on the left panel and Read 2 shown on the right panel. In red are the 14 strains that were sequenced on a date previous to the other 35 strains (grey lines). Two of these strains (SC455 and SC400) that were sequenced on a previous date using PE 100bp reads, in contrast to all other strains.

**Supplementary Figure 3.**
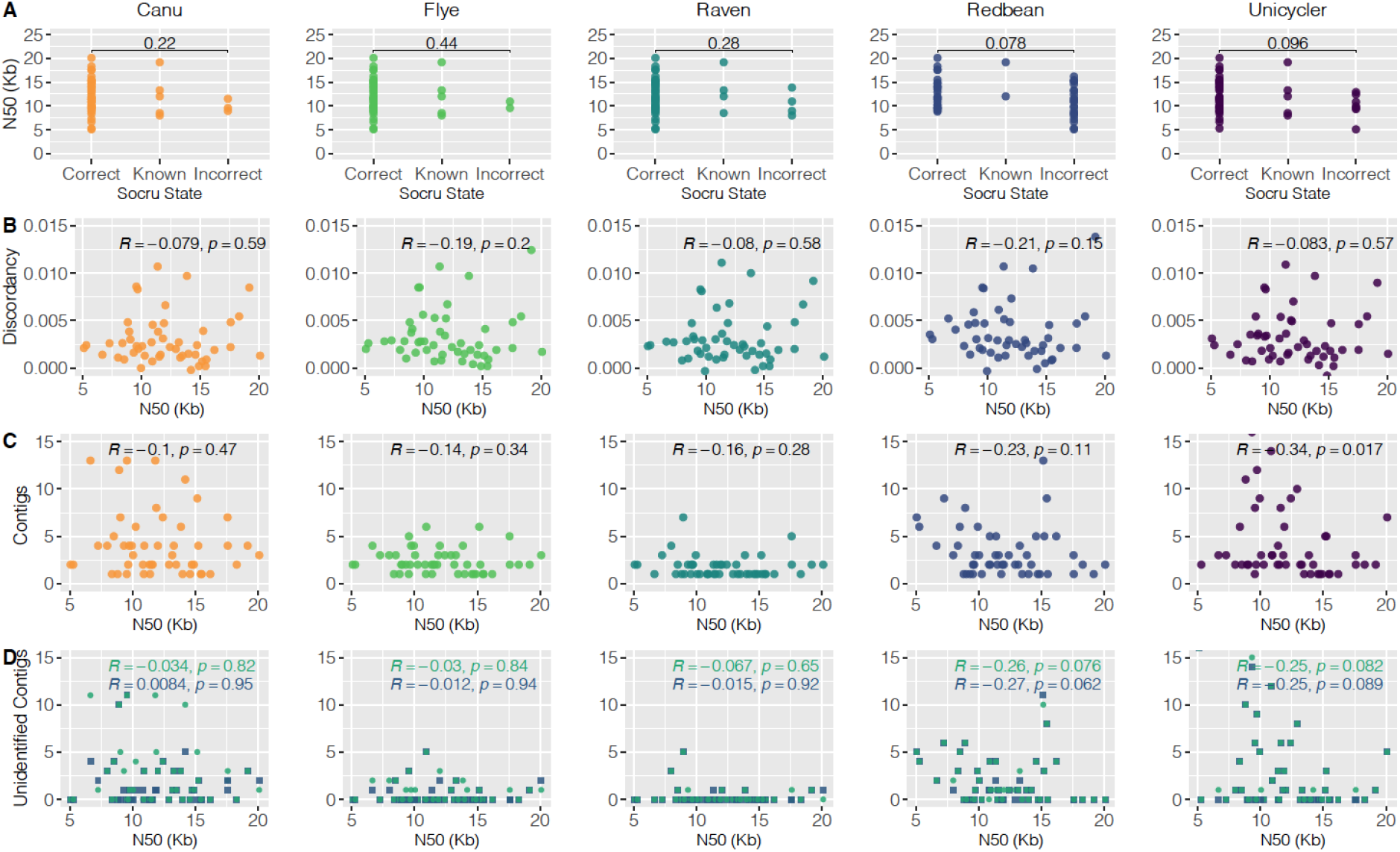
No structural accuracy metrics significantly correlate with read length across strains. **A. rRNA operon ordering vs. read N50.** We tested for significant differences in the read N50 values for correctly or alternatively oriented rRNA operons and incorrectly ordered operons. For all assemblers except Redbean, we did not observe lower N50 values for assemblies with incorrectly oriented operons. **B. Read mapping discordancy vs. read N50.** We found no correlation between the fraction of discordantly mapped reads and N50. Spearman’s rho and corresponding p-values (before correction for multiple comparisons) are shown in each plot. **C. Contig number vs. read N50** and **D. Unidentified contig number vs. read N50.** We found no strong correlation of either contig number or the number of extrachromosomal non-plasmid contigs (“Unidentified contigs”) on read N50. As in **(B)** rho and p-values are shown in each plot.

**Supplementary Figure 4.**
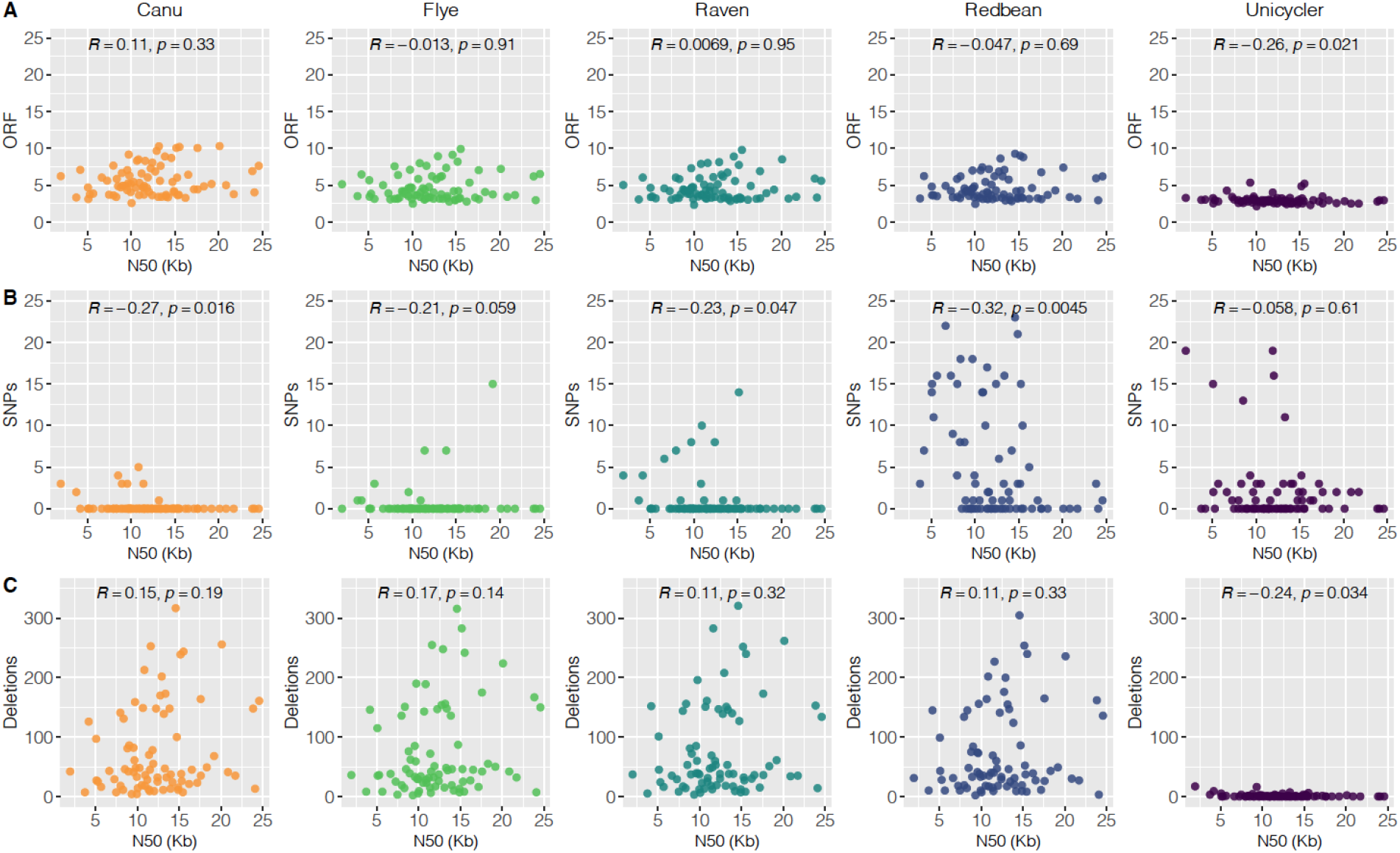
No sequence accuracy metrics significantly correlate with read length across strains. **A. Percentage of short open reading frames. B. Number of SNPs. C. Number of deletions.** SNPs and deletions were called using the breseq pipeline. Spearman’s rho and the corresponding p-values before any correction for multiple comparisons are displayed on each plot.

**Supplementary Figure 5.**
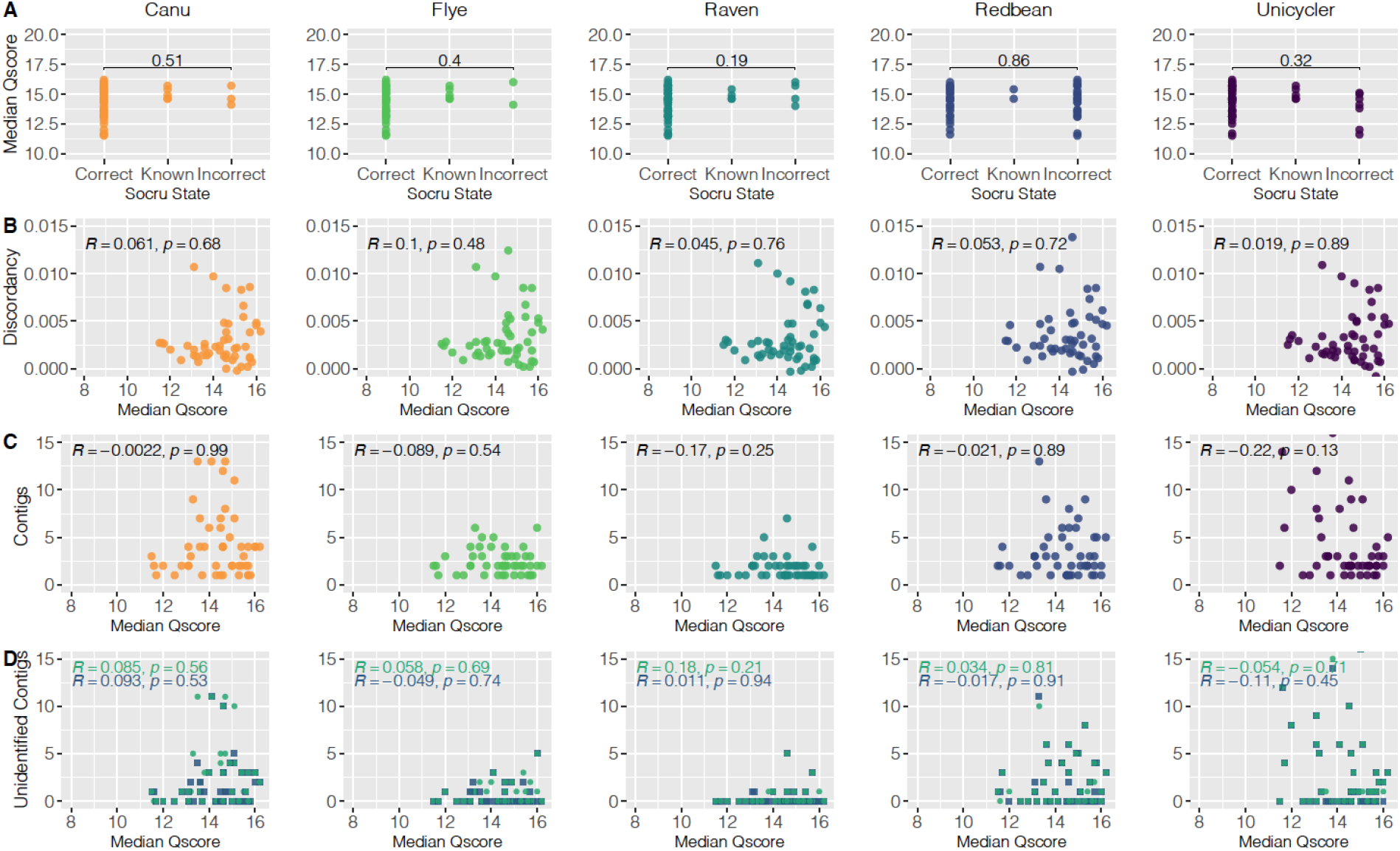
Differences in read quality does not significantly correlate with assembly quality across structural metrics. **A. rRNA operon arrangement. B. Illumina discordancy. C. Total contigs D. Unidentified contig number.** No significant correlation was found across any of the assemblers with the tested metrics. Spearman’s rho and corresponding p-values are displayed on each plot.

**Supplementary Figure 5.**
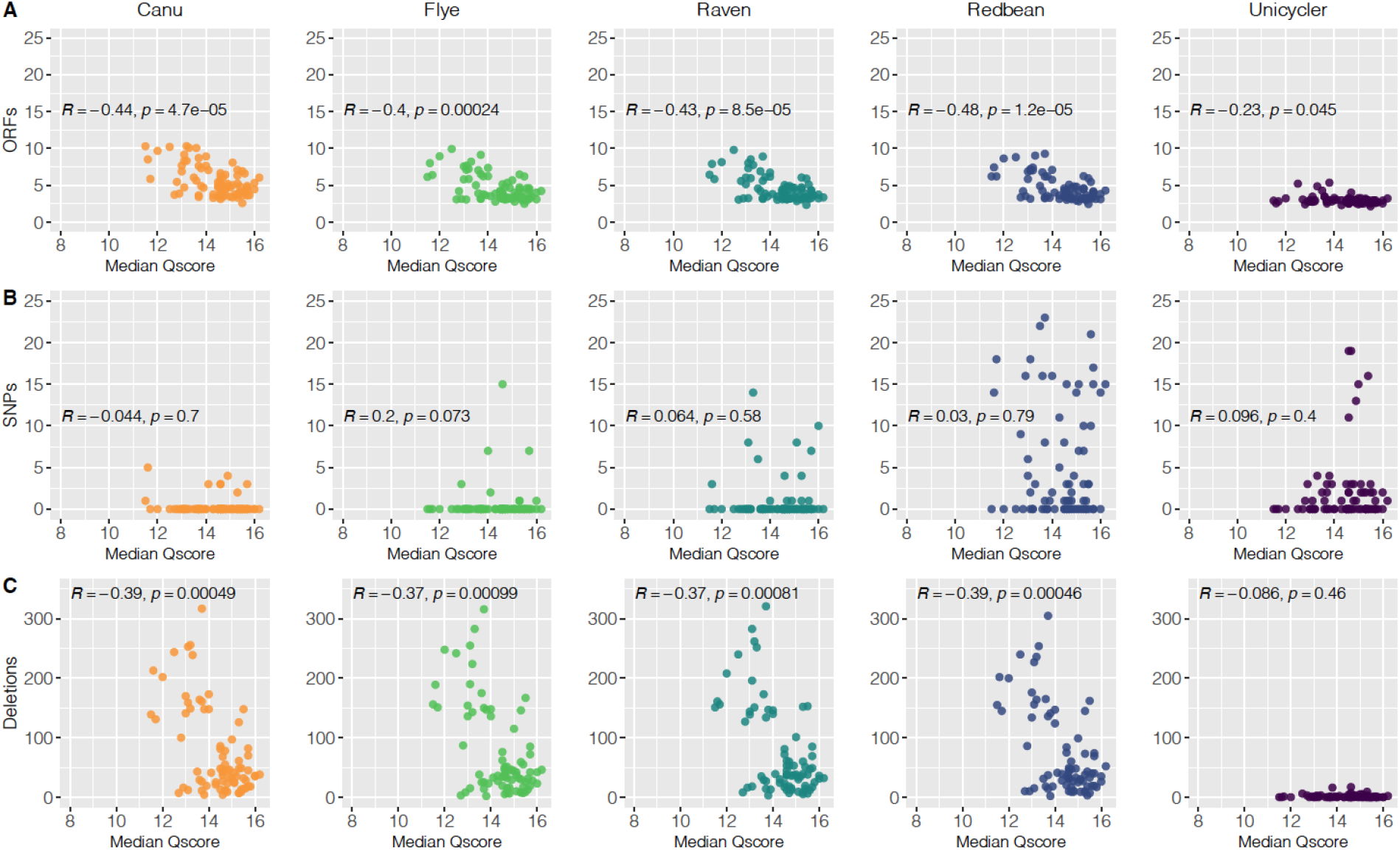
Differences in read quality does not significantly correlate with assembly quality across sequence accuracy metrics. **A. Truncated ORFs B. breseq-called SNPs. C. breseq-called deletions.** No significant correlation was found across any of the assemblers with the tested metrics. Spearman’s rho and corresponding p-values are displayed on each plot.

**Supplementary Figure 7.**
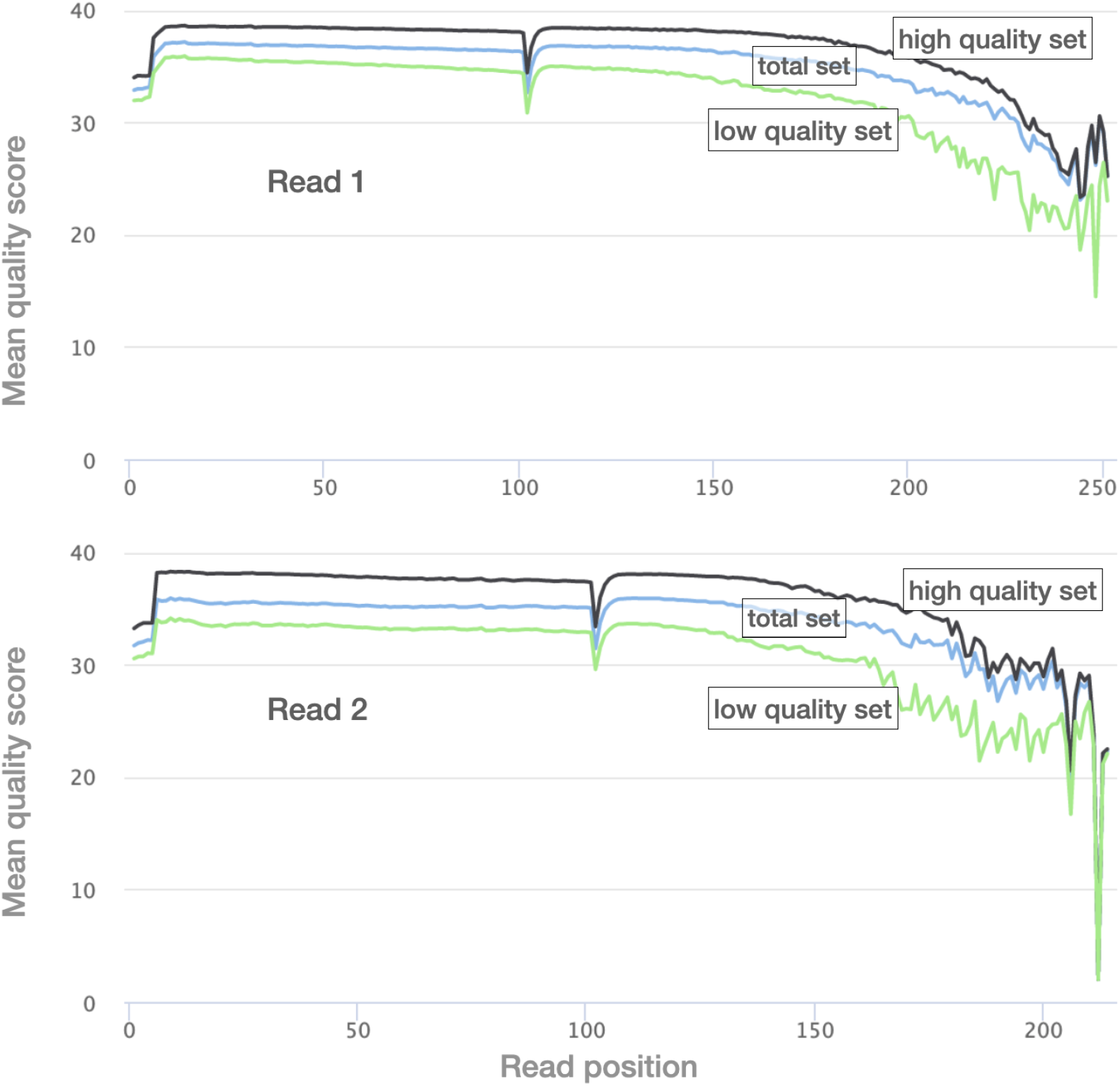
Differences in read quality for “high quality” and “low quality” MG1655 read sets. We used filtlong to divide the MG1655 Illumina data into two read sets, one with high quality reads and the other with low quality reads. Mean quality scores at each position are indicated, calculated using fastp and displayed using multiQC. These different read sets were then used to test the effects of read quality on assembly accuracy.

**Supplementary Figure 8.**
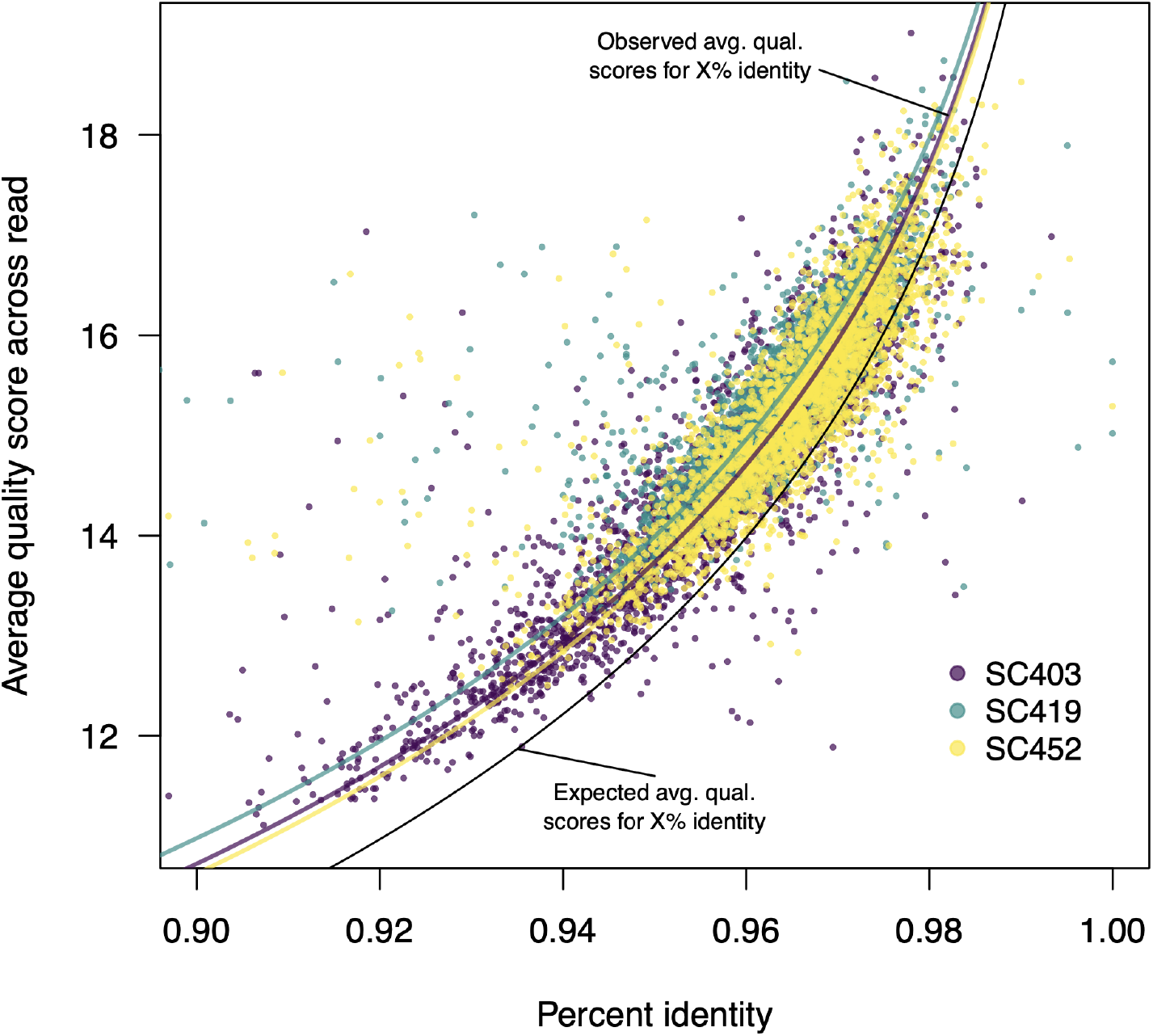
The relationship between read identity and quality score is consistent across strains. We selected three strains (SC403, SC419, and SC452) that differed considerably in the number of truncated ORFs and indels. For each strain, we aligned all reads from a single ONT run to the assembled unicycler genome using minimap2 (Li 2018). We then calculated the percent identity of each read. In the figure above, each point is one read, showing the percent identity of that read and the mean quality score across the read. For clarity, only a subset of the total read numbers are plotted. The black line shows the expected relationship between read identity and average quality. For example we would expect a Q20 read to be 99% accurate and thus have 99% identity to the reference. We generally observed that the mean quality scores were systematically higher than would be expected given the percent identity. For example, a read with 94% identity should, on average, be assigned Q12.2, but we observe most reads with 94% identity assigned as having Q12.8. However, this systematic increase is the same for all strains, showing that Q scores are consistently assigned by the guppy basecaller. The purple, green, and yellow lines indicate a non-linear least squares fits for the relationship between read identity and Q score using an additional parameter that can account for systematic increases or decreases in the relationship between read identity and Q score: Q = −log10(1 - P)*10 + α, in which Q is the quality score, P is the percent identity, and α is a parameter allowing a systematic difference in quality score from what would be expected.

**Supplementary Table 1.**
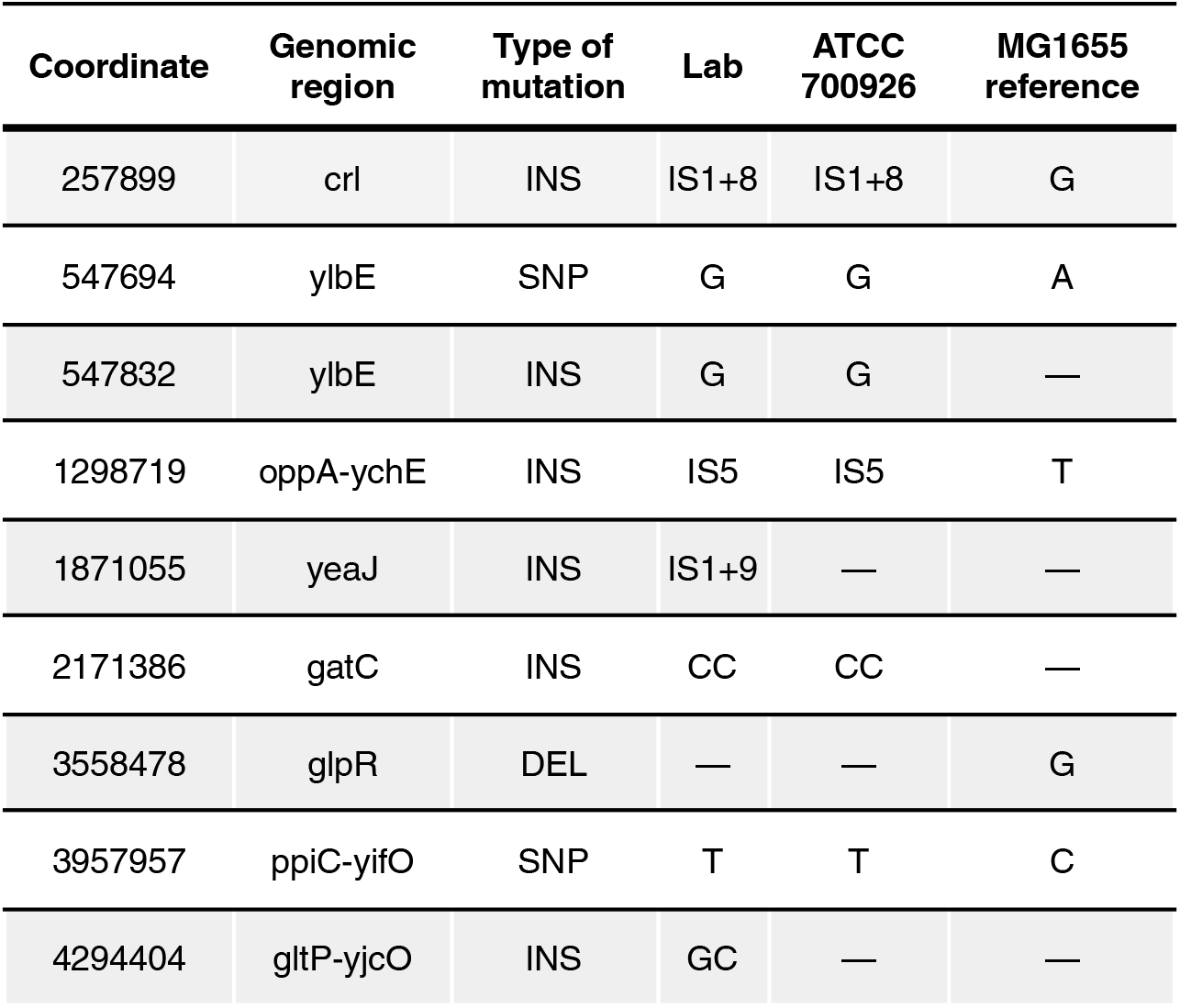
Mutations in the laboratory strain of *E. coli* K12 MG1655. We identified mutations in our laboratory isolate of MG1655 using Illumina reads and the breseq pipeline, which finds only high quality, unambiguous SNPs, indels, and structural rearrangements. The table below shows the differences between our lab strain, ATCC 700926 (distributed by ATCC as the MG1655 genotype), and the MG1655 reference.

**Supplementary Table 2.** Summary of Oxford Nanopore sequencing data for each flow cell. All data were collected using FLO-MIN106 flowcells and either SQK-RBK004 or SQK-LSK109 library preps.

| Run date | Multiplexed samples | Total Mbp | Average mbp per barcode | DNA extraction method | Library |
| --- | --- | --- | --- | --- | --- |
| 2018.06.29 | 6 | 2048 | 341 | Promega | SQK-RBK004 |
| 2018.08.03 | 5 | 2302 | 460 | Promega | SQK-RBK004 |
| 2018.08.10 | 12 | 3919 | 327 | Promega | SQK-RBK004 |
| 2018.08.20 | 4 | 6315 | 1579 | Promega and Phenol | SQK-RBK004 |
| 2018.09.04 | 6 | 8152 | 1359 | Phenol | SQK-RBK004 |
| 2018.09.26 | 10 | 4516 | 452 | Phenol | SQK-RBK004 |
| 2018.09.29 | 8 | 6055 | 757 | Phenol | SQK-RBK004 |
| 2019.02.12 | 10 | 4034 | 403 | Phenol | SQK-RBK004 |
| 2019.02.18 | 11 | 11204 | 1019 | Phenol | SQK-RBK004 |
| 2019.02.20 | 10 | 6077 | 608 | Phenol | SQK-RBK004 |
| 2019.03.20 | 5 | 4145 | 829 | Promega | SQK-RBK004 |
| 2019.03.20 | 8 | 4324 | 540 | Promega | SQK-RBK004 |
| 2021.02.03 | 1 | 4547 | 378 | Promega | SQK-RBK004 |
| 2021.03.12 | 5 | 14850 | 1350 | Promega | SQK-RBK004 |
| 2021.04.07 | 7 | 12703 | 1154 | Promega | SQK-LSK109 |
| 2021.06.04 | 8 | 9461 | 788 | Promega | SQK-LSK109 |
| 2021.07.12 | 8 | 5562 | 427 | Promega | SQK-RBK004 |

**Supplementary Table 3.** Filtered and Unfiltered ONT read quality. ONT datasets were filtered to 500mbp using filtlong.

| Strain | Total reads (thousands) | Total Mbp | Read N50 | Mean read quality score | Reads greater than Q15 | % reads greater than Q15 | Filtered mean read Q score | Filtered reads greater > Q15 | Filtered % greater than Q15 | Filtered read N50 |
| --- | --- | --- | --- | --- | --- | --- | --- | --- | --- | --- |
| SC319 | 108 | 572 | 10613 | 10.7 | 5479 | 5 | 11.4 | 6054 | 8.2 | 10836 |
| SC307 | 128 | 753 | 13855 | 11.3 | 9476 | 7.4 | 13.4 | 10037 | 16.7 | 15159 |
| SC312 | 133 | 770 | 12196 | 9.9 | 3062 | 2.3 | 12 | 3526 | 5.5 | 12913 |
| SC316 | 112 | 817 | 14402 | 10.3 | 3798 | 3.4 | 12.6 | 4430 | 8.2 | 15512 |
| SC364 | 280 | 1734 | 11951 | 10.3 | 9491 | 3.4 | 14.1 | 10740 | 19.7 | 13854 |
| SC368 | 256 | 1637 | 12543 | 10.3 | 6178 | 2.4 | 13.8 | 7412 | 13.9 | 14585 |
| SC375 | 109 | 1118 | 18820 | 10.2 | 3656 | 3.4 | 13.3 | 4297 | 10.9 | 20095 |
| SC386 | 156 | 1344 | 16337 | 10.3 | 5363 | 3.4 | 13.7 | 6328 | 14.1 | 17574 |
| SC392 | 96 | 612 | 14009 | 11.5 | 7596 | 7.9 | 12.8 | 8494 | 14.2 | 14720 |
| SC396 | 245 | 1117 | 9006 | 10.3 | 5744 | 2.3 | 13.1 | 7367 | 9.2 | 9719 |
| SC397 | 113 | 634 | 12518 | 10.2 | 2810 | 2.5 | 11.5 | 3168 | 4.7 | 13167 |
| SC400 | 206 | 966 | 10292 | 10.5 | 6863 | 3.3 | 13.1 | 8452 | 11.4 | 11357 |
| SC402 | 208 | 1208 | 10493 | 9.9 | 3889 | 1.9 | 13.1 | 5001 | 7.6 | 11601 |
| SC403 | 204 | 762 | 7564 | 9.9 | 2906 | 1.4 | 11.8 | 4429 | 4.6 | 8354 |
| SC406 | 416 | 1888 | 7970 | 13.4 | 107636 | 25.9 | 15.8 | 45234 | 84.4 | 10902 |
| SC407 | 601 | 2360 | 7231 | 12.9 | 108432 | 18 | 15.7 | 41525 | 78 | 11395 |
| SC410 | 225 | 1024 | 10591 | 12.9 | 39955 | 17.8 | 15.4 | 31742 | 62.8 | 15439 |
| SC411 | 120 | 789 | 11313 | 13.6 | 29431 | 24.6 | 15 | 28641 | 52.5 | 12411 |
| SC418 | 485 | 1975 | 7463 | 13.5 | 138437 | 28.5 | 16 | 48364 | 89.3 | 10935 |
| SC419 | 193 | 735 | 8384 | 12.1 | 20720 | 10.7 | 14.1 | 21660 | 26.1 | 9556 |
| SC423 | 20 | 155 | 16064 | 13.4 | 5142 | 26 | 13.7 | 5636 | 34.2 | 16195 |
| SC429 | 240 | 1131 | 9096 | 13.4 | 66192 | 27.6 | 15.7 | 44740 | 76.5 | 11500 |
| SC430 | 186 | 474 | 4628 | 13.9 | 58761 | 31.5 | 14.7 | 54388 | 49.3 | 5068 |
| SC431 | 264 | 645 | 4563 | 12.3 | 44945 | 17 | 14 | 46603 | 35.7 | 5274 |
| SC433 | 79 | 681 | 17204 | 13.9 | 28474 | 36.1 | 15.4 | 28004 | 62.3 | 18304 |
| SC434 | 88 | 494 | 10068 | 13.6 | 25734 | 29.3 | 14 | 28242 | 39.3 | 10253 |
| SC441 | 295 | 1043 | 6160 | 13.6 | 84888 | 28.8 | 15.7 | 58850 | 73.5 | 7982 |
| SC443 | 173 | 933 | 10529 | 12.5 | 39064 | 22.5 | 15.4 | 36855 | 62.7 | 12014 |
| SC445 | 27 | 270 | 19143 | 13.7 | 9314 | 33.9 | 14 | 9691 | 41.5 | 19177 |
| SC446 | 35 | 261 | 13212 | 13.8 | 11336 | 32.5 | 14.1 | 12373 | 40.8 | 13288 |
| SC452 | 392 | 2286 | 11753 | 13.6 | 114598 | 29.3 | 16.1 | 32602 | 94 | 17590 |
| SC453 | 133 | 513 | 6911 | 12.6 | 21974 | 16.5 | 13.1 | 24264 | 25 | 7253 |
| SC454 | 275 | 936 | 6855 | 12.2 | 41858 | 15.2 | 15 | 42007 | 48 | 8504 |
| SC455 | 92 | 545 | 11361 | 13.4 | 28341 | 30.8 | 14.4 | 29913 | 43.9 | 11811 |
| SC456 | 28 | 127 | 8608 | 13.7 | 8736 | 31.4 | 14.2 | 9066 | 43.2 | 8935 |
| SC464 | 537 | 2718 | 9395 | 13.4 | 150196 | 28 | 16.2 | 36666 | 95.9 | 15242 |
| SC465 | 109 | 549 | 9016 | 12.7 | 20274 | 18.6 | 13.5 | 23119 | 28.4 | 9315 |
| SC467 | 110 | 519 | 8596 | 13.3 | 30981 | 28.2 | 13.8 | 33226 | 39.4 | 8808 |
| SC469 | 77 | 423 | 9758 | 13.6 | 22484 | 29.2 | 14 | 24802 | 39.7 | 9939 |
| SC475 | 211 | 953 | 8151 | 13.5 | 57510 | 27.2 | 15.5 | 46967 | 67.6 | 10042 |
| SC476 | 365 | 1306 | 7094 | 13 | 82021 | 22.5 | 15.7 | 48656 | 73.5 | 9544 |
| SC477 | 205 | 785 | 7747 | 12.1 | 24497 | 12 | 14.5 | 28549 | 35 | 9030 |
| SC478 | 127 | 981 | 13971 | 13.9 | 45756 | 36.2 | 15.6 | 33812 | 70.9 | 14886 |
| SC479 | 79 | 489 | 11705 | 13.9 | 27559 | 35.1 | 14.2 | 27865 | 43.7 | 11893 |
| SC480 | 77 | 584 | 13064 | 13.8 | 24586 | 32 | 14.8 | 27378 | 48.7 | 13475 |
| SC487 | 110 | 677 | 12907 | 13.5 | 32146 | 29.3 | 15.1 | 31947 | 53.8 | 14199 |
| SC488 | 353 | 1243 | 7044 | 12.9 | 70019 | 19.9 | 15.4 | 41265 | 61.3 | 9652 |
| SC489 | 176 | 530 | 9096 | 12.1 | 21174 | 12.1 | 12.8 | 24685 | 23 | 6644 |
| SC492 | 52 | 410 | 13946 | 13.8 | 16665 | 31.9 | 14.1 | 18436 | 40.8 | 13989 |

## References

Arredondo-Alonso, S., M.R.C. Rogers, J.C. Braat, T.D. Verschuuren, J. Top, J. Corander, R.J.L. Willems, and A.C. Schürch. 2018. Mlplasmids: A User-Friendly Tool to Predict Plasmid- and Chromosome-Derived Sequences for Single Species. Microbial Genomics 4, no. 11. http://www.pubmedcentral.nih.gov/articlerender.fcgi?artid=PMC6321875.

Baltrus, D.A. 2013. Exploring the Costs of Horizontal Gene Transfer. Trends in Ecology & Evolution 28, no. 8 (August): 489–495.

Bertels, F., O.K. Silander, M. Pachkov, P.B. Rainey, and E. van Nimwegen. 2014. Automated Reconstruction of Whole-Genome Phylogenies from Short-Sequence Reads. Molecular Biology and Evolution 31, no. 5 (May): 1077–1088.

Breckell, G., and O.K. Silander. 2020. Complete Genome Sequences of 47 Environmental Isolates of Escherichia Coli. Microbiology Resource Announcements 9, no. 38 (September 17). http://dx.doi.org/10.1128/MRA.00222-20.

Buchfink, B., K. Reuter, and H.-G. Drost. 2021. Sensitive Protein Alignments at Tree-of-Life Scale Using DIAMOND. Nature Methods 18, no. 4 (April): 366–368.

Carattoli, A., E. Zankari, A. García-Fernández, M. Voldby Larsen, O. Lund, L. Villa, F. Møller Aarestrup, and H. Hasman. 2014. In Silico Detection and Typing of Plasmids Using PlasmidFinder and Plasmid Multilocus Sequence Typing. Antimicrobial Agents and Chemotherapy 58, no. 7 (July): 3895–3903.

Chen, S., Y. Zhou, Y. Chen, and J. Gu. 2018. Fastp: An Ultra-Fast All-in-One FASTQ Preprocessor. Bioinformatics 34, no. 17 (September 1): i884–i890.

Deatherage, D.E., and J.E. Barrick. 2014. Identification of Mutations in Laboratory-Evolved Microbes from next-Generation Sequencing Data Using Breseq. Methods in Molecular Biology 1151: 165–188.

De Maio, N., L.P. Shaw, A. Hubbard, S. George, N.D. Sanderson, J. Swann, R. Wick, et al. 2019. Comparison of Long-Read Sequencing Technologies in the Hybrid Assembly of Complex Bacterial Genomes. Microbial Genomics 5, no. 9 (September). http://dx.doi.org/10.1099/mgen.0.000294.

Ewels, P., M. Magnusson, S. Lundin, and M. Käller. 2016. MultiQC: Summarize Analysis Results for Multiple Tools and Samples in a Single Report. Bioinformatics 32, no. 19 (October 1): 3047–3048.

Freddolino, P.L., S. Amini, and S. Tavazoie. 2012. Newly Identified Genetic Variations in Common Escherichia Coli MG1655 Stock Cultures. Journal of Bacteriology 194, no. 2 (January): 303–306.

Goldstein, S., L. Beka, J. Graf, and J.L. Klassen. 2019. Evaluation of Strategies for the Assembly of Diverse Bacterial Genomes Using MinION Long-Read Sequencing. BMC Genomics 20, no. 1 (January 9): 23.

Horesh, G., G.A. Blackwell, G. Tonkin-Hill, J. Corander, E. Heinz, and N.R. Thomson. 2021. A Comprehensive and High-Quality Collection of Escherichia Coli Genomes and Their Genes. Microbial Genomics 7, no. 2 (February). http://dx.doi.org/10.1099/mgen.0.000499.

Hyatt, D., G.-L. Chen, P.F. Locascio, M.L. Land, F.W. Larimer, and L.J. Hauser. 2010. Prodigal: Prokaryotic Gene Recognition and Translation Initiation Site Identification. BMC Bioinformatics 11 (March 8): 119.

Ishii, S., W.B. Ksoll, R.E. Hicks, and M.J. Sadowsky. 2006. Presence and Growth of Naturalized Escherichia Coli in Temperate Soils from Lake Superior Watersheds. Applied and Environmental Microbiology 72, no. 1: 612–621.

Kolmogorov, M., J. Yuan, Y. Lin, and P.A. Pevzner. 2019. Assembly of Long, Error-Prone Reads Using Repeat Graphs. Nature Biotechnology 37, no. 5 (May): 540–546.

Koren, S., and A.M. Phillippy. 2015. ScienceDirect One Chromosome, One Contig: Complete Microbial Genomes from Long-Read Sequencing and Assembly. Current Opinion in Microbiology 23: 110–120.

Koren, S., B.P. Walenz, K. Berlin, J.R. Miller, N.H. Bergman, and A.M. Phillippy. 2017. Canu: Scalable and Accurate Long-Read Assembly via Adaptive *k*-Mer Weighting and Repeat Separation. Genome Research: 1–35.

Kurtz, S., A. Phillippy, A.L. Delcher, M. Smoot, M. Shumway, C. Antonescu, and S.L. Salzberg. 2004. Versatile and Open Software for Comparing Large Genomes. Genome Biology 5, no. 2 (January 30): R12.

Lee, H., T.G. Doak, E. Popodi, P.L. Foster, and H. Tang. 2016. Insertion Sequence-Caused Large-Scale Rearrangements in the Genome of Escherichia Coli. Nucleic Acids Research 44, no. 15 (September): 7109–7119.

Li, H. 2013. Aligning Sequence Reads, Clone Sequences and Assembly Contigs with BWA-MEM (March 16). http://arxiv.org/abs/1303.3997.

Li, H. 2018. Minimap2: Pairwise Alignment for Nucleotide Sequences. Bioinformatics 34, no. 18 (September 15): 3094–3100.

Li, H., B. Handsaker, A. Wysoker, T. Fennell, J. Ruan, N. Homer, G. Marth, G. Abecasis, R. Durbin, and 1000 Genome Project Data Processing Subgroup. 2009. The Sequence Alignment/Map Format and SAMtools. Bioinformatics 25, no. 16 (August 15): 2078–2079.

Long, A., G. Liti, A. Luptak, and O. Tenaillon. 2015. Elucidating the Molecular Architecture of Adaptation via Evolve and Resequence Experiments. Nature Reviews. Genetics 16, no. 10 (October): 567–582.

Marçais, G., J.A. Yorke, and A. Zimin. 2015. QuorUM: An Error Corrector for Illumina Reads. PloS One 10, no. 6 (June 17): e0130821.

Oxford Nanopore Technologies. 2021. Medaka: Sequence Correction Provided by ONT Research. Github. Accessed November 7. https://github.com/nanoporetech/medaka.

Page, A.J., E.V. Ainsworth, and G.C. Langridge. 2020. Socru: Typing of Genome-Level Order and Orientation around Ribosomal Operons in Bacteria. Microbial Genomics 6, no. 7 (July). http://dx.doi.org/10.1099/mgen.0.000396.

Page, A.J., and G. Langridge. 2019. Socru: Typing of Genome Level Order and Orientation in Bacteria. bioRxiv (February 10): 543702.

Quick, J. 2018. Ultra-Long Read Sequencing Protocol for RAD004 Protocol by Josh Quick. http://dx.doi.org/10.17504/protocols.io.mrxc57n.

Ruan, J., and H. Li. 2019. Fast and Accurate Long-Read Assembly with wtdbg2. bioRxiv (January 26): 530972.

Stamatakis, A. 2014. RAxML Version 8: A Tool for Phylogenetic Analysis and Post-Analysis of Large Phylogenies. Bioinformatics 30, no. 9 (May 1): 1312–1313.

Touchon, M., C. Hoede, O. Tenaillon, V. Barbe, S. Baeriswyl, P. Bidet, E. Bingen, et al. 2009. Organised Genome Dynamics in the Escherichia Coli Species Results in Highly Diverse Adaptive Paths. PLoS Genetics 5, no. 1. http://dx.doi.org/10.1371/journal.pgen.1000344.

Vaser, R., and M. Šikić. 2021. Time- and Memory-Efficient Genome Assembly with Raven. Nature Computational Science 1, no. 5 (May 20): 332–336.

Watson, M. 2018. Ideel. Github. Accessed November 9. https://github.com/mw55309/ideel.

Wick, R.R. 2021a. Deepbinner: A Signal-Level Demultiplexer for Oxford Nanopore Reads. Github. Accessed September 4. https://github.com/rrwick/Deepbinner.

Wick, R.R. 2021b. Filtlong: Quality Filtering Tool for Long Reads. Github. Accessed July 27. https://github.com/rrwick/Filtlong.

Wick, R.R., and K.E. Holt. 2019. Benchmarking of Long-Read Assemblers for Prokaryote Whole Genome Sequencing. F1000Research 8 (December 23): 2138.

Wick, R.R., and K.E. Holt. 2021. Polypolish: Short-Read Polishing of Long-Read Bacterial Genome Assemblies. bioRxiv. http://dx.doi.org/10.1101/2021.10.14.464465.

Wick, R.R., L.M. Judd, L.T. Cerdeira, J. Hawkey, G. Méric, B. Vezina, K.L. Wyres, and K.E. Holt. 2021. Trycycler: Consensus Long-Read Assemblies for Bacterial Genomes. Genome Biology 22, no. 1 (September 14): 266.

Wick, R.R., L.M. Judd, C.L. Gorrie, and K.E. Holt. 2017a. Unicycler: Resolving Bacterial Genome Assemblies from Short and Long Sequencing Reads. PLoS Computational Biology 13, no. 6 (June): e1005595.

Wick, R.R., L.M. Judd, C.L. Gorrie, and K.E. Holt. 2017b. Completing Bacterial Genome Assemblies with Multiplex MinION Sequencing. Microbial Genomics 3, no. 10 (October): e000132.

Wick, R.R., M.B. Schultz, J. Zobel, and K.E. Holt. 2015. Bandage: Interactive Visualization of de Novo Genome Assemblies. Bioinformatics 31, no. 20 (October 15): 3350–3352.

